# Reduced GS domain serine/threonine requirements of Fibrodysplasia Ossificans Progressiva mutant type I BMP receptor ACVR1 in the zebrafish

**DOI:** 10.1101/2022.12.01.518722

**Authors:** Robyn S. Allen, William D. Jones, Maya Hale, Bailey N. Warder, Eileen M. Shore, Mary C. Mullins

## Abstract

Fibrodysplasia ossificans progressiva (FOP) is a rare human genetic condition characterized by altered skeletal development and extra-skeletal bone formation. All cases of FOP are caused by mutations in the type I BMP receptor gene *ACVR1* that result in over-activation of the BMP signaling pathway. Activation of the wild-type ACVR1 kinase requires assembly of a tetrameric type I and II BMP receptor complex followed by phosphorylation of the ACVR1 GS domain by type II BMP receptors. Previous studies showed that the FOP mutant ACVR1-R206H requires type II BMP receptors and presumptive GS domain phosphorylation for over-active signaling. Structural modeling of the FOP-ACVR1 mutant kinase domain supports that FOP mutations alter the conformation of the GS domain, but it is unclear how this leads to overactive signaling. Here we show using a developing zebrafish embryo BMP signaling assay that the FOP mutant receptors ACVR1-R206H and -G328R have reduced requirements for GS domain phosphorylation sites to signal compared to wild-type ACVR1. Further, ligand-independent and ligand-dependent signaling through the FOP ACVR1 receptors have distinct GS domain phosphorylation site requirements. Moreover, ACVR1-G328 showed increased GS domain serine/threonine requirements for ligand-independent signaling compared to ACVR1-R206H, whereas it exhibited reduced serine/threonine requirements for ligand-dependent signaling. Remarkably, while ACVR1-R206H does not require the type I BMP receptor partner, Bmpr1, to signal, a ligand-dependent GS domain mutant of ACVR1-R206H could signal independently of Bmpr1 only when Bmp7 ligand was overexpressed. Of note, unlike human ACVR1-R206H, the zebrafish paralog Acvr1l-R203H does not show increased signaling activity. However, in domain-swapping studies the human kinase domain, but not the human GS domain, was sufficient to confer overactive signaling to the Acvr1l-R203H receptor. Together these results reflect the importance of GS domain activation and kinase domain functions in regulating ACVR1 signaling and identify mechanisms of reduced regulatory constraints conferred by FOP mutations.

## Introduction

Fibrodysplasia ossificans progressiva (FOP) is a rare genetic disease characterized by skeletal malformations and extra-skeletal bone formation^1-3^. All cases of FOP are caused by activating mutations in the type I Transforming Growth Factor-β/Bone Morphogenetic Protein (TGF-β/BMP) receptor, ACVR1^4–7^. The most common causative mutation of FOP is a single amino acid change, R206H, within the glycine/serine-rich (GS) domain of ACVR1. Other mutations that cause FOP reside either at alternate sites within the GS domain or in the kinase domain of this receptor^1^. Similar to ACVR1 -R206H, these variant mutations over-activate the BMP signaling pathway and induce phosphorylation of the transcriptional co-activator phospho(p)-Smad1/5^5,7–9^. However, the mechanism by which these mutations over-activate signaling of ACVR1 itself requires further elucidation.

Wild-type ACVR1 signals within a receptor heterotetramer, together with another type I BMP receptor partner and two type II BMP receptors^10–13^. This receptor heterotetramer assembles around the BMP ligand^11,14–17^. Proximity of the receptors within the heterotetramer allows the constitutively active kinase activity of the type II BMP receptor to phosphorylate the type I BMP receptor within its GS domain^18–21^. The GS domain is highly conserved between the multiple type I TGFβ family receptors and across vertebrate and invertebrate species. Within this domain, several serines and a threonine form a Glycine (G)-Serine (S) motif (referred to as the GS loop) for which the domain is named (Table 1). GS domain phosphorylation activates the kinase domain of the type I BMP receptor, allowing it to phosphorylate Smad1/5^22,23^.

**Table 1:**
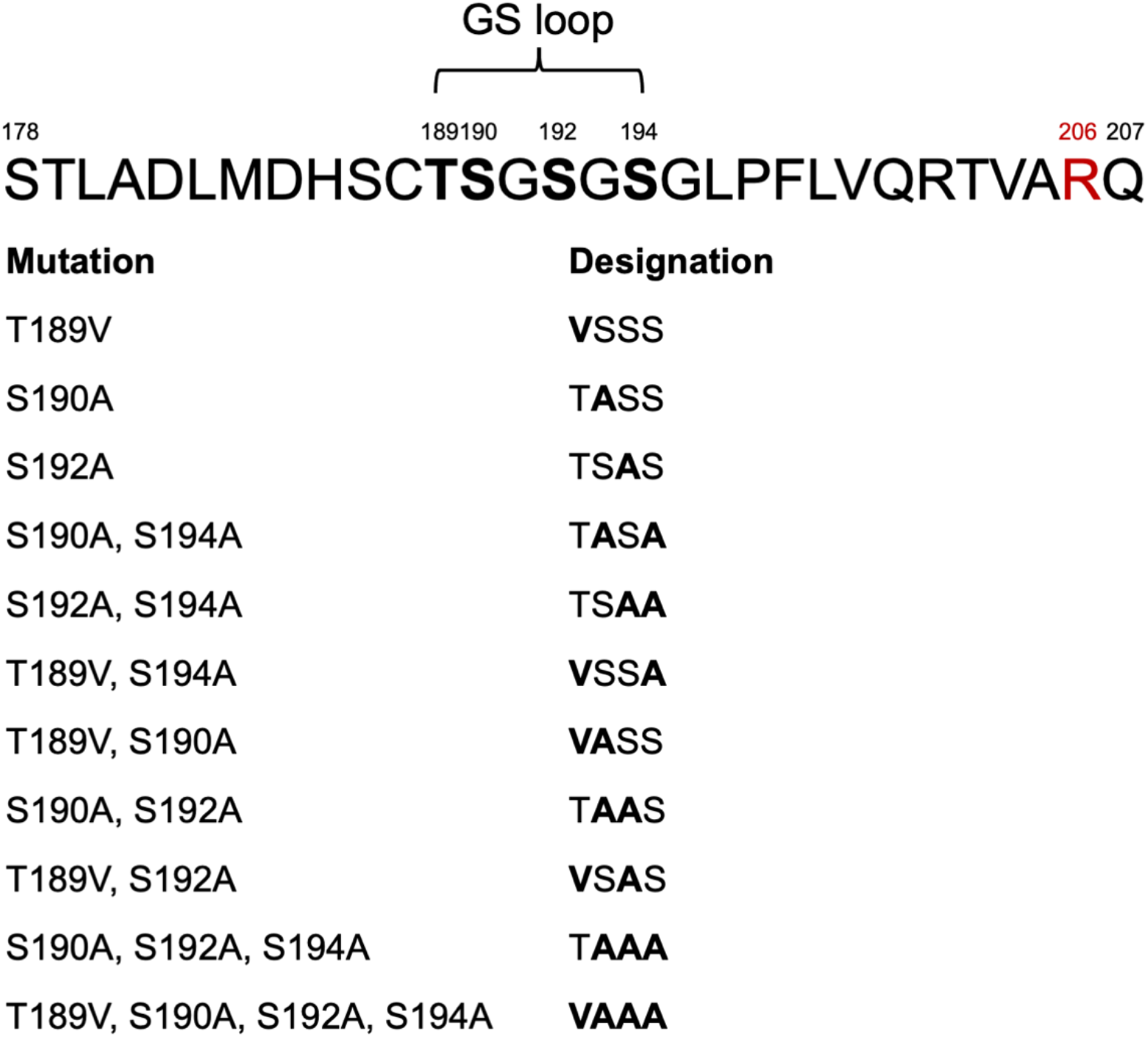
The GS domain of human ACVR1. The GS domain of human ACVR1 spans amino acids 178 to 207. Within this domain are four serine and threonine residues (T189, S190, S192, and S194) that form the GS loop for which the domain is named (Bold, numbered). These residues are phosphorylated by type II BMP receptors. Residue R206 (red, numbered), which is mutated to R206H in FOP, lies within the GS domain. We altered T189, S190, S192, and S194 in the GS loop to unphosphorylatable valine and alanine residues in various combinations as shown.

All the known causative mutations of FOP reside within either the GS domain or the kinase domain of ACVR1. Evaluation of tertiary structures revealed that most of the amino acids altered by ACVR1-FOP mutations lie on the same face of the receptor^1,24^, suggesting that these altered amino acids may over-activate signaling through a similar mechanism. Previous results from our lab and others showed that the FOP mutants ACVR1-R206H and ACVR1-G328R could signal without a ligand binding domain and ACVR1-R206H could signal without its type I BMP receptor partner, Bmpr1, demonstrating that the BMP receptor complex requirements of this mutant receptor are fundamentally altered^5,9,25,26^. However, loss of the type II BMP receptors, BMPR2 and ACVR2a, or the serine/threonine residues within the GS domain abrogate signaling by ACVR1-R206H^27-29^, suggesting that phosphorylation of the GS domain is still required for FOP-ACVR1 to signal.

The zebrafish embryo has been used extensively as a model to assay for BMP signaling by FOP-ACVR1^5,9,25,30^. The zebrafish patterns its dorsal-ventral (DV) embryonic axis through a gradient of BMP signaling activity, a biological mechanism conserved throughout the animal kingdom^12,31,32^. Loss of BMP signaling activity results in expansion of dorsal cell fates at the expense of ventral cell fates, or dorsalization, while overactivation of BMP signaling leads to the opposite effect of ventralization. The use of zebrafish DV patterning as a high throughput screening system for changes in BMP activity led to the discovery of the type I BMP receptor inhibitor, Dorsomorphin^33^. In addition, zebrafish DV patterning was the first *in vivo* model to show that ACVR1-R206H over-activates signaling through pSmad1/5^5,9^. Interestingly, as we report in this study, the synonymous Acvr1l-R203H mutant that is the zebrafish paralog of human ACVR1-R206H does not over-activate signaling. Determining which components of the ACVR1-R206H receptor allow for overactive signaling will provide insight into how this mutation leads to disease and pave the way for an endogenous model of FOP in zebrafish, as well as provide important new insight into the mechanisms that normally regulate BMP receptor activity.

Here we use an *in vivo* zebrafish DV patterning model to evaluate the importance of the GS and kinase domains for overactive signaling by FOP-ACVR1. We show that ACVR1-R206H, a GS domain mutant, and -G328R, a kinase domain mutant (collectively referred to henceforth as FOP-ACVR1), have decreased GS domain phosphorylation site residue requirements compared to WT-ACVR1. In addition, we show for the first time that ligand-independent signaling by ACVR1-R206H and -G328R requires stricter GS phosphorylation site requirements than ligand-dependent signaling by these receptors, with BMP ligand overexpression reducing the requirements for FOP-ACVR1 serine/threonine GS domain residues and for a type I receptor (BMPR1) partner. Moreover, we found that ACVR1-R206H and -G328R differ to each other in their GS residue requirements with ACVR1-G328 exhibiting increased GS residue requirements for ligand-independent signaling but having reduced requirements for ligand-dependent signaling. Further, we found that the human kinase domain, independently of its GS domain, was sufficient to confer overactive signaling to the normally non-overactive zebrafish Acvr1l-R203H. However, swapping only the regulatory αC-helix and A-loop within the kinase domain between human and zebrafish receptors did not alter their activity in the presence of the FOP mutation. Together these data provide further insight into the intricate interactions between ACVR1 domains that facilitate signaling.

## Results

### Ligand-independent signaling of FOP-ACVR1 has stricter GS loop serine/threonine requirements than ligand-dependent signaling of FOP-ACVR1

Previous studies determined that FOP-ACVR1 can signal in the absence of BMP ligand and a ligand binding domain, and that ACVR1-R206H can signal without its type I BMP receptor binding partner BMPR1^25,26^. However, ACVR1-R206H still requires type II BMP receptors and presumptive GS domain phosphorylation^27–29^. Type II BMP receptors activate wild-type (WT) ACVR1 kinase activity via GS domain phosphorylation. Within the GS domain loop of ACVR1 and other type I TGF-β receptors are four serine/threonine residues, T189, S190, S192, and S194 (Table 1), which when phosphorylated cause a conformational change that allows ATP to bind to the kinase domain^20,21^. Loss of two or more of these residues abrogates signaling by the type I TGF-β family receptors, TGFβR1 and ACVR1b, although loss of a single residue had no effect on signaling^20,21,34,35^. In addition, these four residues have been shown to be collectively important for signaling by ACVR1-R206H^28^. Given the evidence that the ACVR1 GS domain R206H mutant receptor may act as a phosphomimetic to decrease the activation threshold of ACVR1-R206H^24,36,37^, we investigated if FOP-ACVR1 has an altered requirement for GS domain phosphorylation sites compared to WT-ACVR1.

To test ACVR1 GS domain activation by type II BMP receptors, we mutated the threonine and serine residues of the ACVR1 GS domain loop to structurally similar, but non-phosphorylatable valine and alanine residues, respectively (Table 1). We then evaluated the signaling activity of these GS mutant receptors using zebrafish DV patterning as an assay for BMP signaling activity (Fig. 1A). Reduced or absent BMP signaling results in expansion of dorsal tissues and loss of ventral tissues (dorsalization, Fig. 1A, C1-C5), while increased BMP signaling results in expansion of ventral tissues and loss of dorsal tissues (ventralization, Fig. 1A, V1-V5). In addition, we previously showed that FOP-ACVR1 has both ligand-independent and ligand-responsive activity^25^. Therefore, we assessed the ability of these GS loop mutant receptors to signal with endogenous BMP ligand, in the absence of BMP ligand, or with excess BMP ligand.

**Figure 1:**
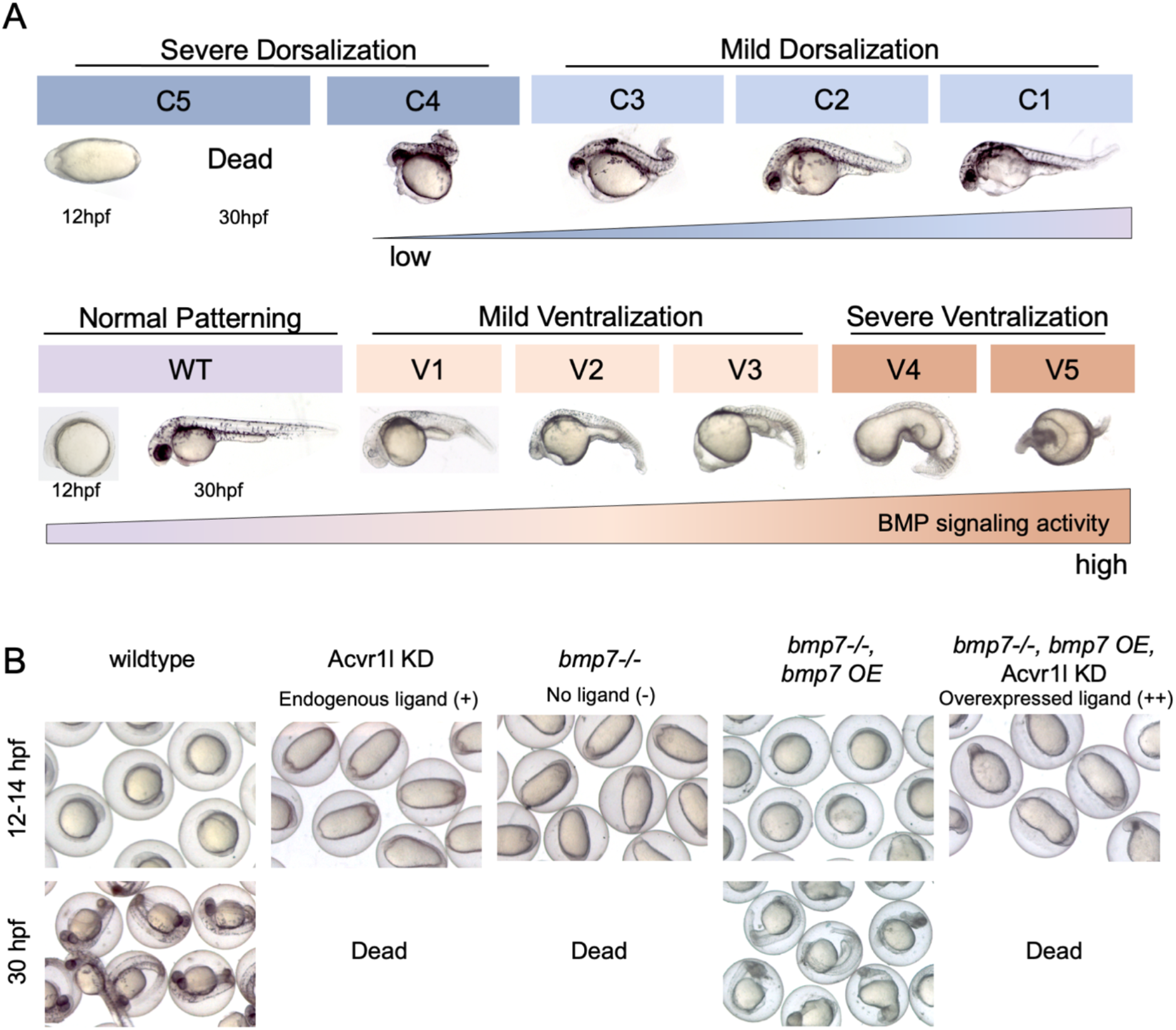
Embryonic zebrafish dorsoventral patterning is a sensitive quantitative assay for BMP signaling activity. **(A)** Dorsoventral patterning phenotypes of zebrafish embryos at 12 hpf (WT and C5) or 28-30 hpf (C4-V5). DV phenotypes range from severe dorsalization (C5-C4, dark blue), mild dorsalization (C3-C1, light blue), wild-type development (WT, violet), mild ventralization (V1-V3, light tan), and severe ventralization (V4-V5, dark tan). C5 embryos are dead by 28 hpf. **(B)** Wild-type and *bmp7*-/- zebrafish embryos with or without Acvr1l KD and/or *bmp7* overexpression (OE) at 12 hpf and 30 hpf. Only live embryos are shown at 30 hpf.

To test the sufficiency of WT- and FOP-ACVR1 GS loop mutants to pattern the zebrafish embryo at a range of Bmp7 levels, endogenous Acvr1l (the zebrafish paralog of human ACVR1) was knocked down and replaced by human GS mutant ACVR1 receptor mRNA injected into embryos with wild-type levels of Bmp7 (endogenous Bmp7 (Bmp7+), no Bmp7 (*bmp7*-/- null mutants (Bmp7-)), or *bmp7*-/- with overexpression (OE) of Bmp7 (Bmp7++) (Table 2). Loss of endogenous Acvr1l in wild-type embryos causes severe dorsalization of the zebrafish embryo to a C5 dorsalized phenotype at 12 hours post fertilization (hpf), characterized by elongation of the embryonic body axis compared to wild-type (Fig. 1A; 1B; 2A, column 2; Supplemental Fig. 1). At 30 hpf, wild-type embryos have a fully patterned body plan, but C5 Acvr1l KD embryos have lysed and died prior to this time point due to radialization of the normally dorsally-restricted somites^38^ (Fig. 1A; 1B; 2A, column 1 and 2; Supplemental Fig. 1). Bmp2/7 heterodimer ligand is the only BMP ligand relevant for patterning the zebrafish embryonic dorsal-ventral axis^11^; as a result, *bmp7*-/- zebrafish embryos display a severely dorsalized C5 phenotype^39^ (Fig. 1B; 2A, column 5; Supplemental Fig. 1). This dorsalized phenotype is rescued to wild-type and ventralized phenotypes by *bmp7* overexpression (Fig 1B; 2A, column 7; Supplemental Fig. 1); but in the absence of Acvr1l, *bmp7* mRNA cannot rescue these embryos because both the receptor and ligand are normally required for patterning (Fig. 1B; 2A, column 8; Supplemental Fig. 1).

**Figure 2:**
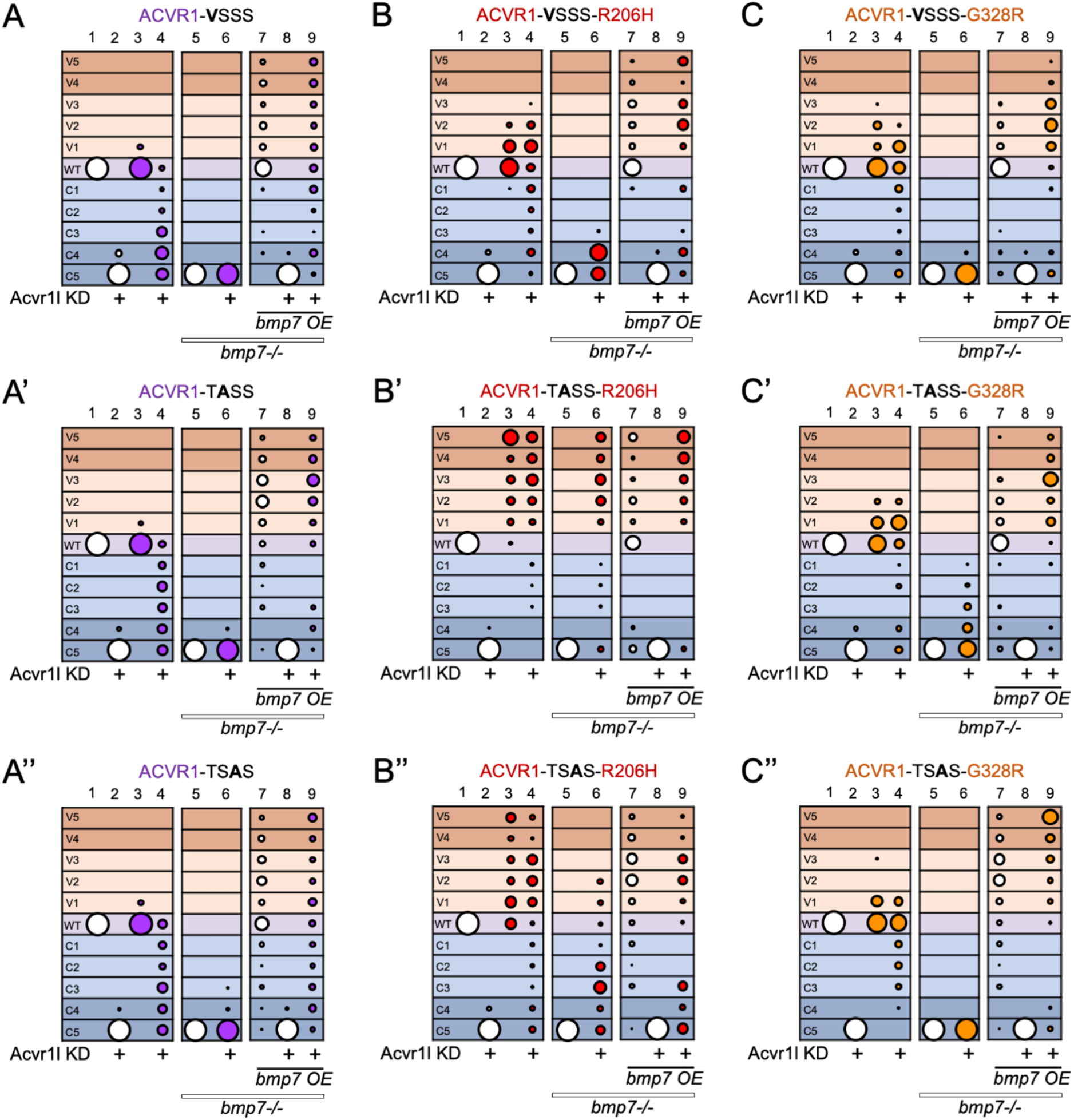
FOP-ACVR1 has differential GS loop S/T requirements for ligandindependent and ligand-responsive signaling. **(A-C)** DV phenotypes of 12-30 hpf wildtype or *bmp7*-/- embryos with or without Acvr1l KD and/or *bmp7* mRNA OE and injected with no receptor mRNA (white circles) or GS loop mutant mRNA of WT-ACVR1 (A, purple), ACVR1-R206H (B, red), or ACVR1-G328R (C, orange). DV phenotypes range from severe dorsalization (C5-C4, dark blue grid background), mild dorsalization (C3-C1, light blue), wild-type development (WT, violet), mild ventralization (V1-V3, light tan), and severe ventralization (V4-V5, dark tan). Bubble plot circle size represents the proportion of embryos with each phenotype at 12-30 hpf. **(A-A”)** *WT-Acvr1* mRNA injected embryos. **(A) V**SSS columns: 1, N=244; 2, N=147; 3, N=71; 4 N=70; 5, N=171; 6, N=891; 7, N=110; 8, N=114; 9, N=99. **(A’)** T**A**SS columns: 1, N=182; 2, N=176; 3, N=123; 4, N=135; 5, N=100; 6, N=52; 7, N=119; 8, N=72; 9, N=45. **(A’’)** TS**A**S columns: 1, N=90; 2, N=74; 3, N=79; 4, N=70; 5, N=144; 6, N=727; 7, N=129; 8, N=77; 9, N=74. **(B-B’’)** *Acvr1-R206H* mRNA injected embryos. **(B) V**SSS columns: 1, N=154; 2, N=150; 3, N=96; 4 N=100; 5, N=176; 6, N=89; 7, N=86; 8, N=82; 9, N=49. **(B’)** T**A**SS columns 1, N=138; 2, N=106; 3, N=89; 4, N=82; 5, N=145; 6, N=95; 7, N=66; 8, N=43; 9, N=107; **(B’’)** TS**A**S columns 1, N=40; 2, N=41; 3, N=86; 4, N=96; 5, N=100; 6, N=51; 7, N=116; 8, N=62; 9, N=37; **(C-C’’)** *Acvr1-G328R* mRNA injected embryos. **(C) V**SSS columns: 1, N=50; 2, N=51; 3, N=92; 4 N=76; 5, N=161; 6, N=72; 7, N=87; 8, N=91; 9, N=72. **(C’)** T**A**SS columns 1, N=137; 2, N=136; 3, N=108; 4, N=74; 5, N=43; 6, N=69; 7, N=73; 8, N=59; 9, N=49. **(C’’)** TS**A**S columns 1, N=63; 2, N=71; 3, N=72; 4, N=72; 5, N=100; 6, N=64; 7, N=116; 8, N=40; 9, N=56. All injections were performed over three or more independent experiments.

**Table 2:**
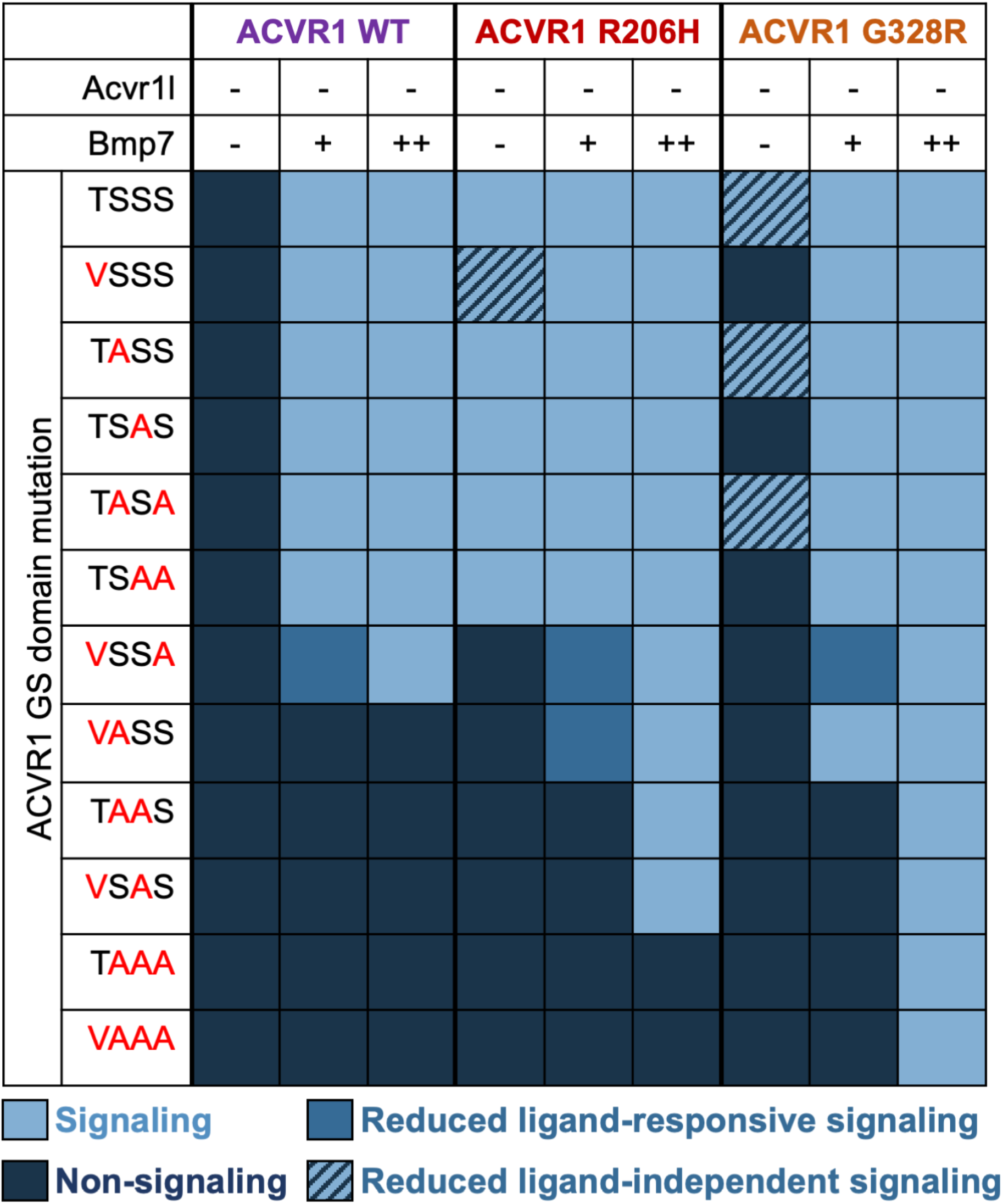
ACVR1 GS loop receptor mutant signaling activity in response to various Bmp7 ligand levels. GS loop mutations (red letters) were introduced into the GS loop of WT-ACVR1, ACVR1-R206H, or ACVR1-G328R constructs. mRNAs of GS loop mutant receptors were assayed for signaling activity in *bmp7* null (-), endogenous *bmp7* (+), or *bmp7* overexpression (++) conditions with knock-down of endogenous Acvr1l (the zebrafish paralog of human ACVR1). Receptors that could rescue more than 5% of Acvr1l KD embryos in these conditions were considered to have signaling activity (light blue, blue, and striped). Receptors that dominant negatively inhibited endogenous Acvr1l, but rescued loss of Acvr1l in endogenous levels of *bmp7* were designated as having reduced ligand-responsive signaling compared to Acvr1l (blue). FOP ACVR1 GS loop mutant receptors that rescued more than 5% of *bmp7* null embryos but did not ventralize these embryos were designated as having reduced ligand-independent signaling (striped). Receptors that could not rescue more than 5% of Acvr1 KD embryos were considered to have no signaling activity (dark blue). Results for ACVR1-TSSS receptors adapted from Allen et al. 2020^26^.

We first tested the ability of WT-ACVR1 to signal with 3 of the 4 phosphorylatable residues in the GS loop. The presence of signaling activity was defined as the ability to rescue more than 5% of C5 embryos to less dorsalized, wild-type or ventralized phenotypes. We examined the ability of these GS domain mutants to signal in different ligand conditions in comparison to WT-ACVR1, ACVR1-R206H, and ACVR1-G328R with intact GS domains (Table 2, TSSS; also previously reported^25^). Normally, WT-ACVR1 overexpression has little effect on signaling in wild-type fish and rescues Acvr1l KD fish to less dorsalized or wild-type phenotypes in the presence of endogenous levels of ligand (Table 2)^25^. WT-ACVR1 cannot rescue *bmp7*-/- embryos, while *bmp7* overexpression ventralizes *bmp7*-/- embryos (Table 2)^25^. Mutation of T189, S190 or S192 to valine or alanine specifically does not alter the rescuing ability of WT-ACVR1 in null mutant, endogenous, or overexpressed Bmp7 (Fig. 2A, A’, A”; Supplemental Fig. 1; Table 2), demonstrating that none of these residues alone is essential for WT-ACVR1 function.

We next tested the ability of the GS domain FOP mutant, ACVR1-R206H, to signal with only three of four phosphorylatable residues. Like ACVR1-TSSS-R206H (wild-type GS loop)^5,25^, ACVR1-**V**SSS-R206H, ACVR1-T**A**SS-R206H and ACVR1-TS**A**S-R206H all ventralized wild-type and Acvr1l KD embryos (Fig. 2B, B’, B”, columns 3 and 4; Supplemental Fig. 2; Table 2). In the absence of Bmp7 ligand, ACVR1-T**A**SS-R206H and ACVR1-TS**A**S-R206H retained the ability to severely ventralize zebrafish embryos (Fig. 2B’, B”, column 6; Supplemental Fig. 2; Table 2). Interestingly, ACVR1-**V**SSS-R206H displayed severely reduced signaling activity in *bmp7*-/- mutants, rescuing them primarily to the strongly dorsalized C4 phenotype (Fig. 2B, column 6; Supplemental Fig. 2). Despite differences in ligand independent signaling, all three mutants still showed enhanced ventralization when Bmp7 ligand was overexpressed (Fig 2B, B’, B’’, column 9; Supplemental Fig. 2; Table 2). These data suggest that T189 may be important for ligand-independent signaling by ACVR1-R206H.

We further investigated the ability of ACVR1-G328R, a kinase domain FOP mutant, to signal with three of four phosphorylatable residues. ACVR1-**V**SSS-G328R, ACVR1-T**A**SS-G328R and ACVR1-TS**A**S-G328R all mildly ventralized wild-type and Acvr1l KD embryos when endogenous ligand was present, similarly to ACVR1-TSSS-G328R^9,25^ (Fig. 2C, C’, C”, columns 3 and 4; Supplemental Fig. 3; Table 2). Previously we showed that ACVR1-TSSS-G328R rescues *bmp7*-/- embryos to less dorsalized phenotypes (Table 2)^25^. Of the single-site GS loop mutants, only ACVR1-T**A**SS-G328R rescued *bmp7*-/- embryos to less dorsalized phenotypes (Fig. 2C’, column 6; Supplemental Fig. 3; Table 2), while ACVR1-**V**SSS-G328R and ACVR1-TS**A**S-G328R rescued less than 5% of embryos to C4 phenotypes in the absence of Bmp7 (Fig. 2C, C”, column 6; Supplemental Fig. 3; Table 2). Doubling or tripling the amount of ACVR1-**V**SSS-G328R or ACVR1-TS**A**S-G328R mRNA failed to increase rescue of *bmp7*-/- mutants above 5%, indicating that their lack of signaling is not due to insufficient receptor expression (Supplemental Fig. 4). All three mutant receptors caused severe ventralization when Bmp7 was overexpressed, demonstrating that the mutant receptors have ligand-responsive functionality (Fig 2C, C’, C”, column 9; Supplemental Fig. 3; Table 2). These results suggest that both T189 and S192 are individually required for ligand-independent signaling by ACVR1-G328R.

Together these data show that no single T/S residue in the GS loop is required for ligand-responsive signaling by any ACVR1 receptors. Although we did not generate single mutants of S194, most double mutants of S194 with T189, S190, or S192, retained activity (Table 2, results reported in next section), indicating that the single loss of S194 would also have normal activity. Interestingly, both FOP-ACVR1 receptors exhibit stricter T/S residue requirements to signal independently of ligand than in responding to ligand, with ACVR1-G328 showing more stringent requirements than ACVR1-R206H (Table 2, compare Bmp7-columns).

### T189 with S190 or S192 is important for ligand-independent signaling by FOP mutant ACVR1

Because ligand mediates assembly of the BMP receptor complex^11,14,40-42^, the presence of ligand facilitates ACVR1 GS domain phosphorylation by type II BMP receptors. However, we and others previously determined that FOP-ACVR1 has the capacity to signal independently of both ligand and an intact ACVR1 ligand binding domain^25,26^. Since T189 appears to be important for ligand-independent, although not ligand-responsive signaling by FOP-ACVR1, we next tested the ability of ACVR1 receptors to signal with only T189 and one other residue.

We first tested the WT-ACVR1 receptor for the sufficiency of two serine/threonine GS domain residues to function in signaling. Like WT-ACVR1-TSSS, ACVR1-TS**AA** and ACVR1-T**A**S**A** rescued loss of endogenous Acvr1l at endogenous ligand levels and ventralized embryos with excess ligand (Fig. 3A, A’, columns 4 and 9; Supplemental Fig. 5; Table 2), demonstrating that two phosphorylatable residues in the GS loop are sufficient for the wild-type ACVR1 receptor to signal. However, ACVR1-T**AA**S inhibited, in a dominant-negative manner, endogenous Acvr1l signaling and failed to rescue loss of endogenous Acvr1l even when Bmp7 was overexpressed (Fig. 3A”, columns 3, 4 and 9; Supplemental Fig. 5; Table 2). Like ACVR1-TS**AA,** ACVR1-T**AA**S was expressed and localized to the cell membrane, although ACVR1-T**AA**S had no apparent signaling activity (Supplemental Fig. 6). Interestingly, ACVR1-**V**SS**A** rescued the loss of endogenous Acvr1l, ventralized embryos overexpressing BMP7, but dominant-negatively dorsalized embryos in the presence of endogenous Acvr1l (Fig. 3A”’, columns 3, 4 and 9; Supplemental Fig. 5; Table 2), suggesting that its signaling activity is reduced compared to wild-type Acvr1l. These data demonstrate that T198 with either S190 or S192 is sufficient for signaling by ACVR1, but S190 and S192 together are not sufficient for normal signaling.

**Figure 3:**
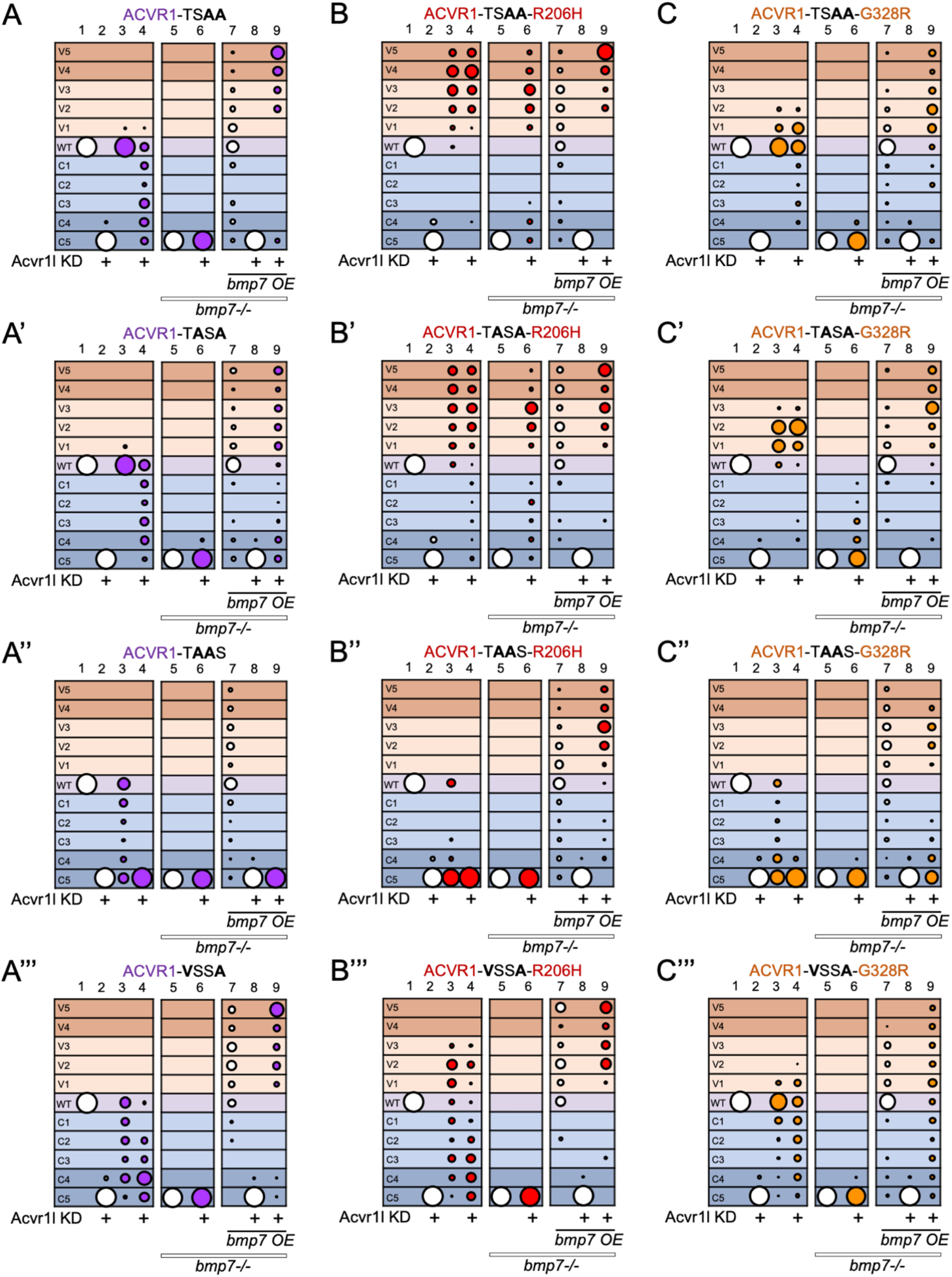
T189 with S190 or S192 is important for ligand-independent signaling by FOP mutant ACVR1. **(A-C)** DV phenotypes of 12-30 hpf wild-type or *bmp7*-/- embryos with or without Acvr1l KD and/or *bmp7* mRNA OE and injected with no receptor mRNA (white circles) or GS loop mutant mRNA of WT-ACVR1 (A, purple), ACVR1-R206H (B, red) or ACVR1-G328R (C, orange). DV phenotypes ranged from severe dorsalization (C5-C4, dark blue grid background), mild dorsalization (C3-C1, light blue), wild-type development (WT, violet), mild ventralization (V1-V3, light tan), and severe ventralization (V4-V5, dark tan). **(A-A”’)** *WT-Acvr1* mRNA injected embryos. **(A)** TS**AA** columns 1, N=287; 2, N=84; 3, N=59; 4, N=60; 5, N=91; 6, N=44; 7, N=38; 8, N=31; 9, N=33. **(A’)** T**A**S**A** columns 1, N=70; 2, N=55; 3, N=79; 4, N=68; 5, N=191; 6, N=156; 7, N=113; 8, N=119; 9, N=131. **(A’’)** T**AA**S columns 1, N=122; 2, N=63; 3, N=68; 4, N=70; 5, N191; 6, N=64; 7, N=64; 8, N=83; 9, N83 **(A”’) V**SS**A** columns 1, N=94; 2, N=76; 3, N=53; 4, N=58; 5, N=159; 6, N=83; 7, N=66; 8, N=57; 9, N=68. **(B-B”’)** *ACVR1-R206H* mRNA injected embryos. **(B)** TS**AA** columns 1, N=104; 2, N=132; 3, N=98; 4, N=106; 5, N=166; 6, N=77; 7, N=76; 8, N=58; 9, N=54. **(B’)** T**A**S**A** columns 1, N=108; 2, N=138; 3, N=99; 4, N=105; 5, N=106; 6, N=83; 7, N=61; 8, N=93; 9, N=60. **(B’’)** T**AA**S columns 1, N=170; 2, N=89; 3, N=68; 4, N=76; 5, N=156; 6, N=122; 7, N=99; 8, N=124; 9, N=57. **(B”’) V**SS**A** columns 1, N=73; 2, N=94; 3, N=73; 4, N=97; 5, N=118; 6, N=84; 7, N=28; 8, N=64; 9, N=67. **(C-C”’)** *ACVR1-G328R* mRNA injected embryos. **(C)** TS**AA** columns 1, N=66; 2, N=83; 3, N=61; 4, N=63; 5, N=123; 6, N=70; 7, N=70; 8, N=61; 9, N=64.**(C’)** T**A**S**A** columns 1, N=158; 2, N=97; 3, N=75; 4, N=68; 5, N=64; 6, N=70; 7, N=35; 8, N=57; 9, N=44. **(C’’)** T**AA**S columns 1, N=148; 2, N=160; 3, N=90; 4, N=82; 5, N=64; 6, N=100; 7, N=71; 8, N=97; 9, N=98. **(C”’) V**SS**A** columns 1, N=76; 2, N=89; 3, N=117; 4, N=127; 5, N=176; 6, N=112; 7, N=115; 8, N=131; 9, N=115. All injections were performed over three or more independent experiments.

We next tested the FOP ACVR1-R206H receptor for the sufficiency of two serine/threonine GS domain residues to function in signaling. ACVR1-TS**AA**-R206H and ACVR1-T**A**S**A**-R206H both rescued Acvr1l KD, and ventralized embryos with and without ligand (Fig. 3B, B’ columns 4, 6 and 9; Supplemental Fig. 7; Table 2), demonstrating that two phosphorylatable residues are sufficient for ligand-independent and ligand-responsive signaling by ACVR1-R206H. Interestingly, ACVR1-T**AA**S-R206H, like ACVR1 -T**AA**S, dorsalized embryos in the presence of endogenous Acvr1l and ligand, but failed to rescue endogenous Acvr1l knockdown (Fig. 3B”, columns 3 and 4; Supplemental Fig. 7; Table 2). These data suggest that ACVR1-T**AA**S-R206H has no signaling activity at endogenous levels of ligand. ACVR1-**V**SS**A**-R206H could both dorsalize and ventralize embryos in the presence of endogenous Acvr1l, indicating that it had variable signaling activity, possibly determined by other yet undefined factors (Fig. 3B”’, column 3 and 4; Supplemental Fig. 7; Table 2). Both ACVR1-T**AA**S-R206H and ACVR1-**V**SS**A**-R206H failed to rescue *bmp7*-/- embryos (Fig. 3B”, B”’ column 6; Supplemental Fig. 7; Table 2), but induced ventralization when Bmp7 was overexpressed (Fig. 3B”, B’” column 9; Supplemental Fig. 7; Table 2). These results suggest that T189 with either S190 or S192 is important for ligand-independent signaling by ACVR1-R206H, but that ligand overexpression relieves the requirement for these specific phosphorylatable residues.

We then tested the FOP ACVR1-G328R receptor to determine if it retains ligand-independent signaling activity with only T189 and either S190 or S192, similarly to ACVR1-R206H. Both ACVR1-TS**AA**-G328R and ACVR1-T**A**S**A**-G328R rescued loss of Acvr1l at endogenous ligand levels (Fig 3C, C’ column 4; Supplemental Fig. 8; Table 2). However, ACVR1-T**AA**S-G328R and ACVR1-**V**SS**A**-G328R dominant negatively inhibited signaling by endogenous Acvr1l, though only ACVR1-**V**SS**A**-G328R still rescued loss of endogenous Acvr1l (Fig 3C”, C”’ column 3 and 4; Supplemental Fig. 8; Table 2). All four of these mutants were ventralizing in the presence of excess ligand (Fig 3C, C’, C”, C”’ column 9; Supplemental Fig. 8; Table 2), but only ACVR1-T**A**S**A**-G328R rescued more than 5% of embryos to less dorsalized phenotypes in the absence of Bmp7 ligand (Fig 3C, C’, C”, C”’ column 6; Supplemental Fig. 8; Table 2). These data indicate that only T189 with S192 is sufficient to allow ligand-independent signaling by ACVR1 -G328R.

Overall, our results show that FOP-ACVR1 requires more phosphorylatable residues in the GS-loop for ligand-independent signaling than ligand-dependent signaling. Further, ACVR1-G328R has stricter GS loop requirements than ACVR1-R206H for ligand-independent signaling, indicating that their mechanisms of conveying overactive signaling may differ.

### Ligand overexpression overrides the requirement for GS loop S/T residues by ACVR1-G328R but not ACVR1-R206H

Since FOP-ACVR1, but not WT-ACVR1, displayed ligand-responsive signaling activity without T189 (Fig 3A”’, B”’, C”’), we next tested the signaling abilities of additional double residue mutants **VA**SS and **V**S**A**S, in addition to GS mutants with one or no phosphorylatable residues. Wild-type ACVR1-**VA**SS, ACVR1-**V**S**A**S, ACVR1-T**AAA** and ACVR1-**VAAA** all dorsalized embryos in the presence of endogenous Acvr1l and failed to rescue loss of endogenous Acvr1l regardless of the amount of Bmp7 present (Fig 4A, A’, A”, A”’; Supplemental Fig. 9; Table 2). All four of these receptors were expressed and localized to the cell membrane, despite having no signaling activity (Supplemental Fig. 10). These data further confirm that T189 with either S190 or S192 is required for signaling by WT-ACVR1.

**Figure 4:**
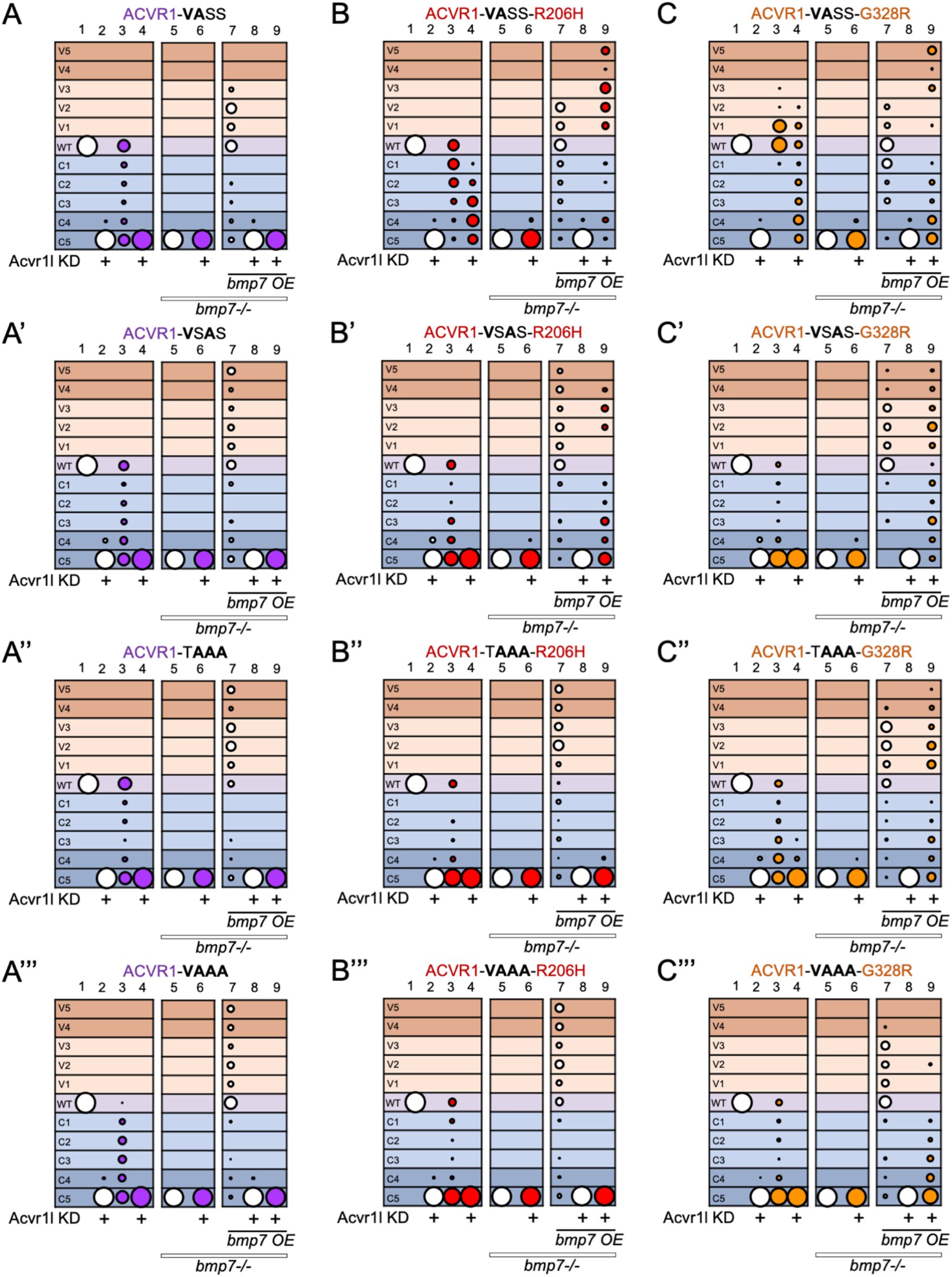
Ligand overexpression overrides the requirement for GS loop S/T residues by ACVR1-G328R but not ACVR1-R206H. **(A-C)** DV phenotypes of 12-30 hpf wild-type or *bmp7*-/- embryos with or without Acvr1l KD and/or *bmp7* mRNA OE and injected with no receptor mRNA (white circles) or GS loop mutant mRNA of WT-ACVR1 (A, purple), ACVR1-R206H (B, red) or ACVR1-G328R (C, orange). DV phenotypes range from severe dorsalization (C5-C4, dark blue grid background), mild dorsalization (C3-C1, light blue), wild-type development (WT, violet), mild ventralization (V1-V3, light tan), and severe ventralization (V4-V5, dark tan). **(A-A”’)** *WT-Acvr1* mRNA injected embryos. **(A) VA**SS columns 1, N=100; 2, N=101; 3, N=60; 4, N=65; 5, N=95; 6, N=92; 7, N=47; 8, N=48; 9, N=92. **(A’) V**S**A**S columns 1, N=194; 2, N=106; 3, N=120; 4, N=117; 5, N=144; 6, N=72; 7, N=158; 8, N=102; 9, N=75. **(A’’)** T**AAA** columns 1, N=126; 2, N=102; 3, N=88; 4, N=81; 5, N=160; 6, N=76; 7, N=65; 8, N=77; 9, N=81. **(A”’) VAAA** columns 1, N=90; 2, N=79; 3, N=95; 4, N=44; 5, N=171; 6, N=77; 7, N=136; 8, N=88; 9, N=63. **(B-B”’)** *Acvr1-R206H* mRNA injected embryos. **(B) VA**SS columns 1, N=287; 2, N=127; 3, N=61; 4, N=65; 5, N=99; 6, N=63; 7, N=31; 8, N=31; 9, N=70. **(B’) V**S**A**S columns 1, N=182; 2, N=74; 3, N=60; 4, N=76; 5, N=194; 6, N=90; 7, N=44; 8, N=67; 9, N=67. **(B’’)** T**AAA** columns 1, N=130; 2, N=146; 3, N=72; 4, N=66; 5, N=160; 6, N=90; 7, N=108; 8, N=65; 9, N=58. **(B”’) VAAA** columns 1, N=90; 2, N=79; 3, N=81; 4, N=75; 5, N=171; 6, N=89; 7, N=87; 8, N=92; 9, N=76. **(C-C”’)** *Acvr1-G328R* mRNA injected embryos. **(C) VA**SS columns 1, N=72; 2, N=96; 3, N=98; 4, N=87; 5, N=118; 6, N=84; 7 N=45; 8, N=37; 9, N=56. **(C’) V**S**A**S columns 1, N=122; 2, N=69; 3, N=68; 4, N=41; 5, N=144; 6, N=42; 7, N=116; 8, N=60; 9, N=70. **(C’’)** T**AAA** columns 1, N=163; 2, N=130; 3, N=93; 4, N=98; 5, N=100; 6, N=99; 7, N=67; 8, N=69; 9, N=79. **(C”’) VAAA** columns 1, N=140; 2, N=114; 3, N=78; 4, N=101; 5, N=76; 6, N=88; 7, N=66; 8, N=40; 9, N=76. All injections were performed over 3 or more independent experiments.

We next tested ACVR1-R206H signaling with these altered GS loop residues. ACVR1-**VA**SS-R206H rescued loss of endogenous Acvr1l, but still dominant negatively inhibited signaling by endogenous Acvr1l (Fig. 4B columns 3 and 4; Supplemental Fig. 11; Table 2), indicating that it has reduced signaling activity compared to endogenous Acvr1l. This receptor failed to rescue *bmp7*-/- mutant embryos but was ventralizing in the presence of Bmp7 overexpression (Fig. 4B columns 6 and 9; Supplemental Fig. 11; Table 2). ACVR1-**V**S**A**S-R206H could not rescue embryos lacking Acvr1l (Fig. 4B’, column 4; Supplemental Fig. 11; Table 2), but in other ways it behaved like ACVR1-**VA**SS-R206H, acting as a dominant negative in a wild-type background (Fig. 4B’ columns 3 and 4; Supplemental Fig. 11; Table 2), unable to rescue patterning in the absence of Bmp7 (Fig. 4B’ column 6; 11), and induced ventralization in response to excess Bmp7 (Fig. B’ column 9; Supplemental Fig. 11). Neither ACVR1-T**AAA**-R206H nor ACVR1-**VAAA**-R206H rescued loss of endogenous Acvr1l at any ligand level (Fig. 4B”, B”’ columns 4, 6 and 9; Supplemental Fig. 11; Table 2). We confirmed that both these receptors were expressed and localized to the cell membrane (Supplemental Fig. 10). These data show that ACVR1-R206H requires phosphorylatable residues within the GS loop and that T189 is not sufficient for signaling even when ligand is overexpressed.

Finally, we investigated the ability of ACVR1-G328R to signal without T189. Unlike ACVR1-**VA**SS-R206H, ACVR1-**VA**SS-G328R displayed little dominant-negative inhibition of endogenous Acvr1l (Fig. 4C, column 3; Supplemental Fig. 12; Table 2). ACVR1-**VA**SS-G328R mildly ventralized zebrafish embryos in endogenous and excess Bmp7 (Fig. 4C, columns 4 and 9; Supplemental Fig. 12) but did not rescue loss of Acvr1l in the absence of ligand (Fig 4C column 6; Supplemental Fig. 12). ACVR1-**V**S**A**S-G328R, and, remarkably, ACVR1-T**AAA**-G328R and ACVR1-**VAAA**-G328R, while having no signaling activity when ligand is absent or at endogenous levels (Fig 4C’, C’’, C”’ columns 4 and 6; Supplemental Fig. 12; Table 2), were able to rescue Acvr1l knockdown embryos in the presence of excess Bmp7 (Fig 4C’, C”, C”’ columns 9; Supplemental Fig. 12; Table 2). These receptors localized normally to the cell membrane (Supplemental Fig. 10). These data show that ligand overexpression allows ACVR1-G328R but not ACVR1-R206H to signal in the absence of any phosphorylatable GS loop residues and supports different mechanisms of overactive signaling by ACVR1-R206H and ACVR1-G328R. The results of the receptor activity assays in Figures 2–4 are summarized in Table 2.

### ACVR1-VASS-R206H but not ACVR1-TSAA-R206H requires Bmp7 overexpression to signal when BMPR1 is deficient

In the zebrafish, Bmp2/7 ligand heterodimers are necessary for Acvr1l to form complexes with Bmpr1^11,13^. The requirement for ligand by certain GS domain FOP-ACVR1 mutants suggests that these mutants also require receptor complex assembly. To test if there is a differential type I receptor requirement between our ligand-independent and ligand-dependent ACVR1-R206H GS loop mutants, we generated zebrafish embryos deficient for all type I BMP receptors by intercrossing *bmpr1aa*+/-;*bmpr1ab*-/- zebrafish and injecting their embryos with morpholinos (MOs) against *bmpr1ba, bmpr1bb* and *acvr1l* [referred to as type I knockdown (KD) embryos]. We then injected these embryos with mRNAs encoding the ligand-independent GS loop mutant, ACVR1-R206H-TS**AA**, or the ligand-dependent GS loop mutant, ACVR1-R206H-**VA**SS, with or without *bmp7* mRNA overexpression. We scored embryo phenotypes without knowledge of their genotypes, and then subsequently genotyped the embryos.

Bmpr1 (encoded by the *bmpr1aa, bmpr1ab, bmpr1ba* and *bmpr1bb* genes in zebrafish) is normally required for BMP signaling to pattern the DV axis of the developing zebrafish embryo^11,13,25^. As previously described, *bmpr1aa*+/-;*bmpr1ab*-/- (referred to here as *bmpr1a*+/-) zebrafish embryos develop normally, but *bmpr1aa*-/-; *bmpr1ab*-/- (referred to as *bmpr1a*-/-) embryos are dorsalized to a C4 phenotype (Fig 5A; 5E, column 1 and 7). Type I KD embryos, lacking all type I BMP receptors, are severely dorsalized to a C5 phenotype (Fig 5B; 5E, column 3 and 9). *bmp7* overexpression ventralizes wild-type and *bmpr1a*+/- embryos, but cannot rescue *bmpr1a*-/- or type I KD embryos, showing that normally Acvr1l requires Bmpr1a to respond to ligand (Fig 5C; 5D; 5E, column 2, 4, 8 and 10)^25^. ACVR1-TS**AA**-R206H over-activated signaling and ventralized type I KD embryos with or without *bmp7* overexpression (Fig 5E, columns 11 and 12). In contrast, ACVR1-R206H-**VA**SS modestly rescued Acvr1l KD embryos with endogenous levels of Bmp7 when Bmpr1a was present (Fig 5F, column 5), but rescue was abrogated in type I KD embryos lacking Bmpr1 (Fig 5F, column 11). However, when Bmp7 was overexpressed ACVR1-R206H-**VA**SS was able to ventralize type I KD embryos (Fig 5F, column 12). These data show that ligand-dependent ACVR1-**VA**SS-R206H signaling requires Bmpr1 at endogenous ligand levels, but ligand overexpression overcomes this requirement. In contrast, ACVR1-TS**AA**-R206H can signal at endogenous ligand levels even when Bmpr1 is deficient, showing that T189 and S190 convey enhanced signaling activity compared to S192 and S194 in FOP-ACVR1-R206H.

**Figure 5:**
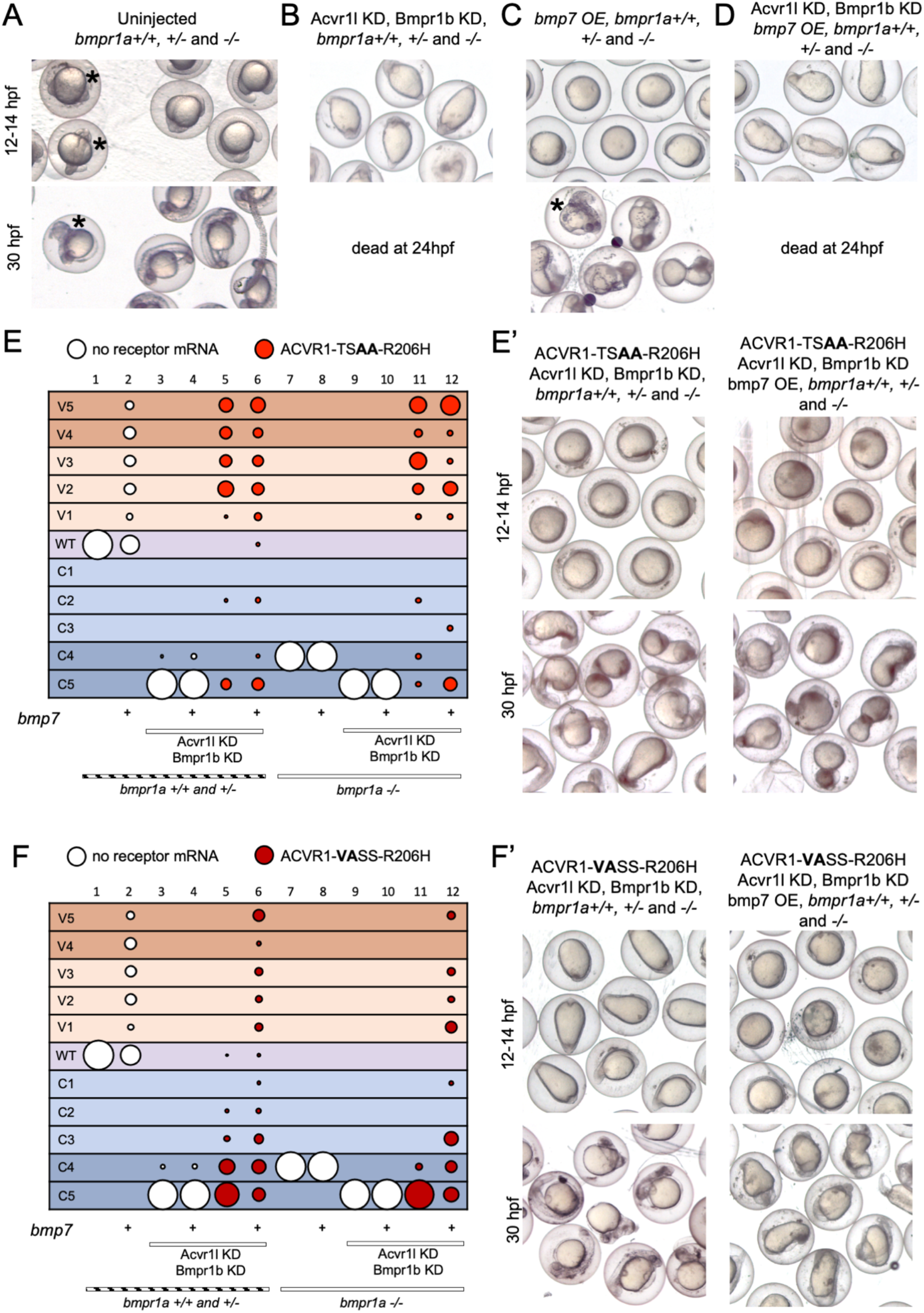
ACVR1-R206H-VASS but not ACVR1-R206H-TSAA requires Bmp7 overexpression to signal in the absence of Bmpr1. **A-D)** DV phenotypes of 12-30 hpf *bmpr1a*+/+, +/- and -/- embryos, that are uninjected or injected with *bmpr1b* morpholino, *acvr1* morpholino, and/or *bmp7* mRNA. Asterisk indicates examples of non-rescued C4 *bmpr1a*-/- embryos (A and C). **(E)** *Acvr1-TS**AA**-R206H* (ligand-independent GS loop mutant) mRNA injected embryos with or without *bmp7* mRNA. Columns: 1, N=36; 2, N=80; 3, N=83; 4, N=77; 5, N=60; 6, N=54; 7, N=46; 8, N=16; 9, N=24; 10, N=32; 11, N=28; 12, N=26. **(E’)** Representative 12 and 30 hpf phenotypes. **(F)** *Acvr1-**VA**SS-R206H* (ligand-dependent GS mutant) mRNA injected embryos with or without *bmp7* mRNA. Columns: 1, N=76; 2, N=91; 3, N=106; 4, N=95; 5, N=118; 6, N=127; 7, N=58; 8, N=27; 9, N=34; 10, N=39; 11, N=35; 12, N=40. **(F’)** Representative 12 and 30 hpf phenotypes. All images show a mix of *bmpr1a*+/+, +/- and -/- embryos. Injections were performed over 3 independent experiments. Only live embryos are shown at 30 hpf (Fig 5B, D).

### The human ACVR1 kinase domain, but not the human GS domain, is sufficient to facilitate overactive signaling by zebrafish Acvr1l-R203H

Previous analysis of the 3-D structure of ACVR1 showed that all known FOP mutations reside on the same face of the receptor, near the GS domain^1^. The proximity of GS domain FOP mutants and some kinase domain FOP mutants to the GS loop suggests that the mutations act to destabilize the GS loop and the normally inhibitory conformation of the receptor in the absence of ligand^24,37^. The intracellular domains of human ACVR1 and zebrafish Acvr1l are highly conserved, with over 91% identity at the amino acid sequence between the GS domains and 85% identity between the kinase domains (Fig. 6A), suggesting that the FOP mutations would behave similarly in zebrafish Acvr1l and human ACVR1.

**Figure 6:**
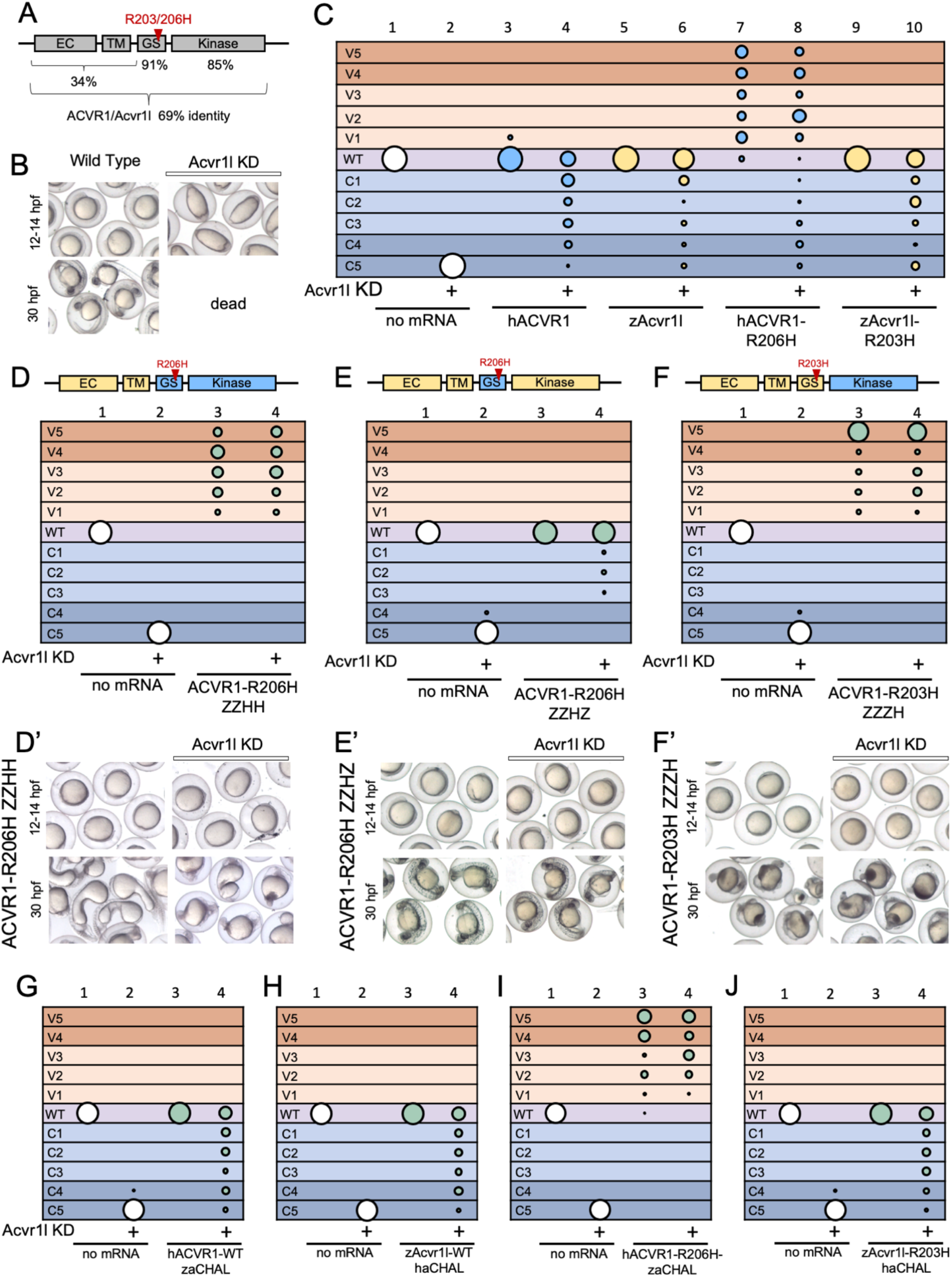
The human ACVR1 kinase domain is sufficient to confer overactive signaling by zebrafish Acvr1l-R203H. **(A)** Amino acid identity between human ACVR1 and zebrafish Acvr1l extracellular (EC), transmembrane (TM), GS, and Kinase domains. Schematic not to scale. **(B)** Representative 12 and 30 hpf phenotypes of wild-type embryos with or without Acvr1l KD. **(C-J)** DV phenotypes of 12-30 hpf wild-type embryos with *acvr1l* KD injected with no mRNA (white) or mRNAs for human (h, H) ACVR1 (blue), zebrafish (z, Z) Acvr1l (yellow), or chimeric zebrafish-human ACVR1 (green). **(C)** Embryos with or without Acvr1l KD injected with zebrafish *acvr1l*, zebrafish *acvr1l-R203H*, human *Acvr1* or human *Acvr1-R206H* mRNAs. Columns: 1, N=202; 2, N=215; 3, N=72; 4, N=61; 5, N=86; 6, N=87; 7, N=83; 8, N=93; 9, N=51; 10, N=51. **(D-F)** Schematics above plots show domains of each chimera from human ACVR1 (blue) or zebrafish Acvr1l (yellow) **(D)** Embryos with or without Acvr1l KD injected with the *Acvr1-R206H-ZZHH* chimeric mRNA (zebrafish extracellular and transmembrane domains with human GS and kinase domains. Columns: 1, N=33; 2, N=56; 3, N=47; 4 N=41. **(D’)** Representative 12 and 30 hpf phenotypes. **(E)** Embryos with or without Acvr1l KD injected with the *Acvr1l-R206H-ZZHZ* chimeric mRNA (zebrafish extracellular, transmembrane and kinase domains, and human GS domain). Columns: 1, N=80; 2, N=73; 3, N=100; 4, N=91. **(E’)** Representative 12 and 30 hpf phenotypes. **(F)** Embryos with or without Acvr1l KD injected with the *Acvr1l-R203H-ZZZH* chimeric mRNA (zebrafish extracellular, transmembrane and GS domains, and human kinase domain). Columns: 1, N=80; 2, N=73; 3, N=76; 4, N=74. **(F’)** Representative 12 and 30 hpf phenotypes. **(G)** Embryos with or without Acvr1l KD injected with human WT-ACVR1 with zebrafish αC helix and A-loop (aCHAL) (L250I, R275K, T356M). Columns: 1, N=138; 2, N=121; 3, N=87; 4, N=94. **(H)** Embryos with or without Acvr1l KD injected with zebrafish WT-Acvr1l-haCHAL (I247L, K372R, T362M). Columns: 1, N=98; 2, N=99; 3, N=66; 4, N=61. **(I)** Embryos with or without Acvr1l KD injected with human ACVR1-R206H-zaCHAL (L250I, R375K, M365T. Columns: 1, N=142; 2, N=135; 3, N=74; 4, N=90. **(J)** Embryos with or without Acvr1l KD injected with zebrafish Acvr1-R203H-haCHAL (I247L, K372R, T362M). Columns: 1, N=193; 2, N=102; 3, N=86; 4, N=135. Injections were performed over three or more independent experiments. Only live embryos are shown at 30 hpf (Fig 6B).

To determine if zebrafish Acvr1l-R203H, the paralog to human ACVR1-R206H, can over-activate signaling, we injected zebrafish *acvr1l-R203H* or human *Acvr1-R206H* mRNAs into wild-type zebrafish embryos with or without endogenous Acvr1l KD (Fig. 6B). Surprisingly, zebrafish Acvr1l-R203H behaved similarly to WT-Acvr1l and WT human ACVR1 and was unlike human ACVR1-R206H. Acvr1l-R203H did not perturb normal patterning in wild-type zebrafish embryos, unlike the ventralization induced by ACVR1-R206H (Fig. 6C, columns 3, 5, 7, and 9) and rescued Acvr1l KD fish to less dorsalized or wild-type phenotypes (Fig. 6C, columns 4, 6, and 10; Supplemental Fig. 13), in contrast to ACVR1-R206H, which ventralized them (Fig. 6C, column 8). Because the R206H mutation lies within the GS domain, these data suggest there may be a functional difference between the human and zebrafish GS domain that facilitates overactive signaling by the mutant receptor from one species but not the other.

To investigate if the intracellular domains of the human receptor could confer overactivity to the zebrafish Acvr1l receptor, we generated a human-zebrafish chimeric receptor in which the zebrafish extracellular and transmembrane domains are fused to the human GS and kinase intracellular domains (ACVR1l-R206H-ZZHH; zebrafish amino acids 1-174 (analogous to human 1-177), human amino acids 178-509). We then tested the ability of this chimeric receptor to rescue the C5 dorsalized phenotype of Acvr1l KD embryos or over activate signaling to cause ventralization. We found the zebrafish-human chimera ACVR1l-R206H-ZZHH ventralized embryos regardless of the presence of endogenous Acvr1l (Fig 6D and D’). These results show that the human ACVR1 intracellular domain is responsible for facilitating overactivity by the R206H mutation to the zebrafish extracellular and transmembrane domains.

We next tested which part of the human ACVR1 intracellular domain, the GS and/or the kinase domain, could confer overactivity to the zebrafish Acvr1l receptor. We generated chimeric receptors in which either the human GS or kinase domain replaced the respective domain of the zebrafish Acvr1l receptor. We found that ACVR1l-R206H-ZZHZ (zebrafish amino acids 1-174 and 205-506 (analogous to human 1-177 and 208-509), human amino acids 178-207), which contains the human GS domain and zebrafish extracellular, transmembrane and kinase domains, rescued loss of endogenous Acvr1l, but did not ventralize zebrafish embryos (Fig. 6E and E’). Halving or doubling the amount of *Acvr1l-R206H-ZZHZ* mRNA injected into WT zebrafish embryos did not alter the ability of the receptor to rescue Acvr1l KD or allow it to over-activate signaling (Supplemental Fig. 14). These results show that the differences between the human and zebrafish GS domains where the R206H mutation resides are not responsible for the overactive signaling of R206H. Interestingly, however, ACVR1-R203H-ZZZH (zebrafish amino acids 1-204 (analogous to human 1-207), human amino acids 208-509), which contains the zebrafish extracellular, transmembrane and GS domains, with the human kinase domain, did over-activate signaling, severely ventralizing embryos regardless of the presence of endogenous Acvr1l (Fig. 6F and F’). Both ACVR1-R206H-ZZHZ and ACVR1-R203H-ZZZH were expressed and localized to the cell membrane (Supplemental Fig. 15). Altogether, these results demonstrate that the human kinase domain is sufficient to confer ventralizing overactivity to the zebrafish Acvr1l-R203H receptor and suggest that an interaction between the GS and kinase domains is critical for the pathogenesis of FOP.

Finally, we investigated if two regulatory structures within the kinase domain, the αC-helix and the A-loop, might explain the difference between human ACVR1-R206H and zebrafish Acvr1-R203H. The αC-helix and the A-loop, both help mediate opening of the ATP binding pocket within the ACVR1 kinase domain^37,43^. Three amino acids within or adjacent to the αC-helix and A-loop differ between human ACVR1 and zebrafish Acvr1l, L250/I247, R375/K372 and M365/T362. Of note, a variant FOP mutation, R375P, lies within these structures^1^. We generated receptors with the αC-helix and the A-loop (aCHAL) residues swapped between human(h) and zebrafish(z) and tested their ability to rescue loss of endogenous Acvr1l and over-activate signaling. Swapping the sequences of the αC-helix and A-loop between human WT-ACVR1 and zebrafish WT-Acvr1l had no effect on their normal signaling activity (Fig. 6G and H; Supplemental Fig. 16). Similarly, swapping the αC-helix and A-loop between human FOP-ACVR1-R206H and zebrafish Acvr1l-R203H had no effect on their signaling activities. hACVR1-R206H-zaCHAL retained its ability to over-activate BMP signaling and ventralize zebrafish embryos (Fig 6I, Supplemental Fig. 16). zAcvr1l-R203H-haCHAL behaved like wild-type zAcvr1l and zAcvr1l-R203H, rescuing loss of endogenous Acvr1l without displaying overactive signaling (Fig 6J; Supplemental Fig 16). These results suggest that the function of the Cα-helix and A-loop structures is conserved between human and zebrafish and does not contribute to their difference in FOP-R206H mutant activity. Overall, these data begin to identify the molecular mechanism through which ACVR1-R206H gains overactive signaling that is not imparted to zebrafish Acvr1-R203H, and suggest possible uncharacterized regulatory structures within the kinase domain that are playing a key role.

## Discussion

Using an *in vivo* zebrafish development model as a quantitative assay for BMP signaling activity, we are the first to show that ligand-responsive and ligand-independent signaling through FOP-mutant ACVR1-R206H and -G328R receptors have distinct GS loop serine/threonine (S/T) residue requirements. Specific mutations to the GS loop residues that are normally phosphorylated in activated type I BMP receptors generated non-signaling, ligand-responsive, or ligand-independent receptors. Interestingly, a ligand-dependent FOP-ACVR1 GS loop mutant also re-acquired dependence on Bmpr1 to signal in response to endogenous levels of ligand. Finally, we show that the human ACVR1 kinase domain, but not the GS domain, is sufficient to confer overactive signaling activity to zebrafish Acvr1l-R203H. These data demonstrate that the GS and kinase domains have highly coordinated functional activities and this coordination is integral to the pathogenesis of FOP.

Our results show that for both wild-type ACVR1 and FOP-ACVR1, specific combinations of two of the four S/T residues within the GS loop that are phosphorylated by the BMP type II receptors are sufficient to signal in response to endogenous ligand levels. The wild-type ACVR1 GS loop mutants that retained the ability to signal required T189 in combination with either S190 or S192. Similarly, GS mutant FOP-ACVR1 receptors that retained T189 with either S190 or S192 did not have altered signaling compared to intact FOP-ACVR1-TSSS receptors. Of note, compared to these receptors, the double S/T residue mutants ACVR1-**V**SS**A**, ACVR1-**V**SS**A**-R206H, and ACVR1-**V**SS**A**-G328R, displayed reduced ligand-dependent signaling activity, rescuing embryos deficient in endogenous Acvr1l to less dorsalized, wildtype, or even ventralized phenotypes, but dominant-negatively inhibiting endogenous Acvr1l as shown by their dorsalizing effects on wild-type zebrafish embryos (Fig 3B”’, C”’, Fig 4B). The double S/T residue mutants ACVR1-**VA**SS, ACVR1-**VA**SS-R206H, and ACVR1-**VA**SS-G328R all behaved distinctly to each other. ACVR1-**VA**SS could no longer signal, even when BMP7 ligand was overexpressed, whereas ACVR1-**VA**SS-R206H behaved like ACVR1-**V**SS**A**-R206H exhibiting reduced activity, and ACVR1-**VA**SS-G328R retained signaling activity resembling intact ACVR1-TSSS-G328R (Fig 4C). These data demonstrate that FOP mutations reduce the GS domain phosphorylation site residue requirement of ACVR1, and that in the presence of endogenous ligand ACVR1-G328R displayed a less specific phosphorylation residue requirement for signaling than ACVR1-R206H.

Previously, we and others reported that unlike WT-ACVR1, ACVR1-R206H and ACVR1-G328R signal in the absence of ligand^25,26^. Strikingly, we show here that some FOP-ACVR1 GS loop mutants retained the ability to signal in the absence of Bmp7 ligand, while others could only signal in the presence of Bmp7 ligand, and still others required Bmp7 ligand overexpression to signal (summarized in Table 2). Five of the ACVR1-R206H and two of the ACVR1-G328R GS loop mutants retained the ability to signal in the absence of ligand. ACVR1-**V**SSS-R206H, ACVR1-T**A**SS-R206H, ACVR1-TS**A**S-R206H, ACVR1-T**A**S**A**-R206H, ACVR1-TS**AA**-R206H, ACVR1-T**A**SS-G328R, and ACVR1-T**A**S**A**-G328R all showed some level of ligand-independent signaling activity. Notably, all the ligand-independent ACVR1-R206H mutants ventralized embryos in the absence of ligand, with the exception of ACVR1-**V**SSS-R206H. ACVR1-**V**SSS-R206H consistently could rescue *bmp7*-/- embryos only to a severely dorsalized C4 phenotype, indicating that in the absence of T189, ligand-independent signaling by ACVR1-R206H is severely impaired. Together, these data show that T189 with either S190 or S192 is important for ligand-independent signaling by the FOP mutant, ACVR1-R206H. The FOP kinase domain mutant, ACVR1-G328R, only exhibited modest ligand-independent activity when both T189 and S192 were retained, showing that ACVR1-G328R has a stricter GS domain residue requirement than ACVR1-R206H for ligand-independent signaling. This is consistent with data showing that ACVR1-G328R with an intact GS domain exhibits lower ligand-independent activity than ACVR1-R206H^8,9^. This lends further support to R206H and G328R mutations conferring over-active signaling by different mechanisms.

Six FOP GS loop receptor mutants could signal only when ligand was overexpressed, including two ACVR1-G328R mutants that lack all or nearly all the S/T residues of the GS loop. Bmp7 overexpression allowed ACVR1-T**AA**S-R206H, ACVR1-**V**S**A**S-R206H, ACVR1-T**AA**S-G328R, and ACVR1-**V**S**A**S-G328R mutants to signal. Excess ligand was not sufficient to allow the non-signaling wild-type ACVR1 GS mutants to signal, suggesting that the FOP mutant residues indeed act to reduce the GS domain phosphorylation requirements for receptor activation. While ACVR1-**TAAA**-R206H and ACVR1-**VAAA**-R206H failed to signal even when ligand was overexpressed, remarkably, ACVR1-**TAAA**-G328R and ACVR1-**VAAA**-G328R acquired signaling activity in the presence of Bmp7 overexpression. These data suggest that not only does the G328R mutation abrogate the need of the GS loop S/T residues in the presence of ligand overexpression, but that ligand may play a role in BMP receptor complex signaling distinct from initiating GS loop phosphorylation, possibly by causing a conformational change or relief of inhibition that has not been previously described. Alternatively, ACVR1-**VAAA**-G328R may be phosphorylated at sites other than those within the GS loop to activate signaling. Previous experiments in cell culture showed that residue T203 alone was sufficient to allow ACVR1-**VAAA**-R206H to signal when co-expressed with type II BMP receptors^44^, suggesting that in conditions of excess receptor components (i.e., ligand or type II receptors), phosphorylatable residues outside the GS loop are sufficient to allow kinase activation. The type II receptors, ACVR2a and BMPR2 have been shown to be important for signaling by ACVR1-R206H^27,29^. Of note, ACVR2a is more responsive to both BMP7 ligand (used in this study) and Activin ligand (a novel activator of FOP-ACVR1), while BMPR2 responds more efficiently to BMP4 ligand^27,45-47^. As a result, Bmp7 ligand overexpression may recruit excess ACVR2a for signaling and reflect the importance of this receptor for FOP-ACVR1. These data suggest that excess ligand or type II BMP receptor may increase the likelihood of a phosphorylation event, which in turn may compensate for loss of S/T residues within the GS loop.

A ligand-independent and a ligand-responsive GS loop mutant showed differential requirements for the type I BMP receptor partner, Bmpr1. Previously, we showed that ACVR1-R206H can signal in the absence of Bmpr1, the obligate type I BMP receptor partner of WT-ACVR1 in the zebrafish embryo^11,25^. Like ACVR1-R206H, ACVR1-TS**AA**-R206H ventralized zebrafish embryos in the absence of Bmpr1, showing that it retains Bmpr1 independent signaling activity. Conversely, ACVR1-**VA**SS-R206H, a GS loop mutant that requires ligand to signal, also required the type I BMP receptor partner Bmpr1 to signal, demonstrating that these GS domain mutations cause it to re-acquire endogenous BMP signaling complex requirements (Fig 5). Interestingly, however, Bmp7 overexpression allowed ACVR1-**VA**SS-R206H, but not endogenous WT-Acvr1l, to signal independently of Bmpr1 (Fig 5). Increased ligand may increase the interactions between ACVR1 and type II receptors alone or with novel type I and II receptor complexes. Overall, these data further support that when T189 and S190 are phosphorylatable, ACVR1 can signal independently of its normal tetrameric complex.

Collectively these results further elucidate the roles of BMP ligand-receptor complex components in FOP-ACVR1 signaling. The ability of FOP-ACVR1 to signal in the absence of ligand and Bmpr1 suggests the ability to signal in the absence of a BMP ligand-receptor complex. However, in a genetic mouse model of FOP, WT-ACVR1 overexpression lessens the severity of the phenotype^48^, presumably through competition for BMP ligand-receptor components with ACVR1-R206H. These data support that FOP-ACVR1 signals more efficiently with its canonical BMP ligand-receptor complex components than without^48^. Our results and others support that phosphorylation of the GS domain is required for activation of ACVR1-R206H, suggesting an important role for phosphorylation by type II BMP receptors^27-29^. Previous studies, however, showed that ACVR1-R206H and the constitutively active ACVR1-Q207D receptor could signal with kinase dead type II receptors^29^, supporting that while type II receptors are required for signaling, their kinase activity is dispensable. Alternative sources of phosphorylation may come from transphosphorylation by other type I receptors or by autophosphorylation by ACVR1 itself. T189 may be the GS loop phosphorylation site that is more amenable to these alternative phosphorylation mechanisms. The importance of T189 for ligand-independent signaling by FOP-ACVR1 indicates T189 as the site most accessible for phosphorylation or the site with the greatest impact on receptor activity without ligand and potentially complex assembly.

Changes in the interaction between ACVR1-R206H and the ACVR1 regulator FKBP12 have been proposed to explain the increased signaling activity observed in ACVR1-R206H. In the absence of ligand, the FOP ACVR1-R206H mutant receptor displays reduced interactions with the type I BMP receptor inhibitor FKBP12, which is normally only displaced upon ligand binding and complex assembly^6,37,49,50^. Reduced binding of FKBP12 may allow certain GS loop residues to be more readily phosphorylated in the absence of ligand-mediated complex assembly.

It is important to note that although valine and alanine are highly structurally similar to threonine and serine, respectively, they differ in polarity: threonine and serine are polar while valine and alanine are non-polar. We cannot exclude the possibility that changes in the charge of amino acids within the GS domain may lead to unexpected changes in protein conformation and affect receptor activation in unexpected ways. However, in support of a lack of protein conformation changes is that ligand-dependent signaling of wild-type ACVR1 is normal when any one of the Ser/Thr residues is altered to Ala/Val and for the two double Ser substitutions to Ala tested.

Finally, our data show that a specific interaction between the GS and kinase domains is critical for the pathogenesis of FOP in the human ACVR1 but this interaction does not occur in the zebrafish paralog Acvr1l. We determined that Acvr1l-R203H, the zebrafish paralog to the human ACVR1-R206H, does not over-activate BMP signaling in the zebrafish embryo (Fig 6). Swapping the human and zebrafish GS domains did not confer increased signaling to zAcvr1l-“R203H”, indicating that increased signaling is not solely dependent on changes within the GS domain. However, when the kinase domain of zAcvr1l-R203H was replaced by the human ACVR1 kinase domain, the chimeric zAcvr1l-R203H receptor became over-activating. These data indicate that important but undefined interactions between the GS and kinase domain regulate over-activity of the receptor. We additionally determined that the αC helix and A-loop, two kinase domain structures that regulate the ATP binding pocket^37,43^, are not responsible for this species-specific difference in signaling activity. An important future direction in resolving this novel mechanism of receptor regulation will be to determine which human residues are required to impart overactive signaling to the zebrafish receptor to better understand how these receptors are regulated in signaling. Taken together these data elucidate the function of multiple domains within ACVR1 and give us new insight into how these receptors are regulated both normally and in disease.

## Methods

### Study design

All injection experiments into wildtype or *bmp7*-/- embryos were performed in a minimum of 50 embryos over 2 separate experiments. All injection experiments into *bmpr1a*+/-;*bmpr1b*-/- incrossed animals were performed in a minimum of 150 embryos to obtain at least 30 double mutant injected animals over 2 separate experiments. No statistical sample estimations were used in designing this study.

Evaluation of results: In all experiments, rescue of BMP signaling was defined as more than 5% of injected embryos patterning to a C4-V5 phenotype, with a C5 phenotype representing lack of BMP signaling. This threshold was chosen as the cut off for signaling because overexpression of WT-ACVR1 consistently does not rescue more than 5% of *bmp7*-/- embryos to a C4 phenotype or above. Due to lack of an appropriate statistical comparison tool and challenges accounting for differences in mRNA activity, this study refrained from comparing relative rescue between mutant receptors. Each receptor was independently evaluated for its ability to perturb wildtype development or rescue severely dorsalized C5 embryos to less dorsalized, wildtype, or ventralized phenotypes. The one exception was our claim that ACVR1-VSSS-R206H appears to have severely reduced ligand-independent signaling activity due to its consistent rescue of *bmp7*-/- to a C4 phenotype.

### Zebrafish

All procedures involving animals were performed with approval by the University of Pennsylvania IACUC. Zebrafish husbandry was carried out according to institutional and national ethical and animal welfare guidelines. Adult zebrafish were housed in schools of 20-50 fish in a 13hr light/11hr dark day/night cycle. Wild-type Tübingen zebrafish were used for all experiments involving wild-type fish. The following established mutant lines were used in this study: *bmp7a^sb1aub^* (*bmp7*-/-)^39^ and *bmpr1aa^p3/+^*; *bmpr1ab^sa0028^* (*bmpr1aa*+/-;*bmpr1ab*-/-)^25^. *bmp7a^sb1aub^* were maintained as homozygous mutant stocks by injecting 1-cell stage embryos with *bmp7a* mRNA to rescue the C5 dorsalized phenotype. *bmpr1a* mutant fish stocks were maintained as *bmpr1aa*+/-;*bmpr1ab*-/- heterozygotes. Adults were incrossed and *bmpr1aa*+/-;*bmpr1ab*-/- offspring were identified by genotyping. Double homozygous null mutants (*bmpr1aa*-/-;*bmpr1ab*-/-) are embryonic lethal. Embryos at 1-48 hpf were maintained in 28-32°C E3 solution for all experiments. Sex was not accounted for as zebrafish sex determination does not take place until juvenile stages of development^51^.

| **Zebrafish genotype** | **Source/ Catalog#** | **Strain** | **Providing laboratory** |
| --- | --- | --- | --- |
| Wild-type Tübingen (TU) | NA | TU | NA |
| *bmp7a*<sup>sb1aub</sup> | NA | TU | Mullins lab;<br>University of<br>Pennsylvania<sup>39</sup> |
| *bmpr1ab*<sup>sa0028</sup> | ZIRC/ZL2118 | TU | NA |
| *bmpr1aa*<sup>p3/+</sup> | NA | TU | Mullins lab,<br>University of<br>Pennsylvania<sup>25</sup> |

### Genotyping

DNA samples for genotyping were obtained from adult zebrafish fin clips or whole zebrafish embryos collected in 100% methanol. DNA was isolated using HotShot DNA extraction. *bmpr1aa* genotyping was performed using the KASPar genotyping system^52^. Proprietary primers were used for genotyping as previously described (LGC Biosearch technologies)^25^. *bmpr1a* +/+(*bmpr1aa*+/+;*bmpr1ab*-/-), *bmpr1a*+/-(*bmpr1aa*+/-;*bmpr1ab*-/-) and *bmpr1a*-/- (*bmpr1aa*-/-;*bmpr1ab*-/-)_*embryos* were generated from *bmpr1aa*+/-; *bmpr1ab*-/- incrosses. Following injection and blind phenotyping, embryos were genotyped as described above.

### Site-directed mutagenesis

GS loop mutant receptors were generated using site directed mutagenesis. Plasmids were mutated sequentially; one or two -basepairs were mutated in each reaction. pCS2+ backbones containing ACVR1 or an FOP-ACVR1 mutant were PCR amplified with the mutant primer using Phusion high fidelity polymerase (NEB, M0530). Mutated plasmids were transformed into Top10 chemically competent cells and DNA from individual colonies was submitted for Sanger sequencing. The full coding sequence of ACVR1 or FOP-ACVR1 was sequenced to ensure that no unwanted mutations were incorporated during the mutagenesis process.

### PCR and In-Fusion cloning

Chimeric human-zebrafish ACVR1 receptors were generated using PCR amplification and In-Fusion cloning. Primers were designed to amplify a desired segment of the receptor with an overhang corresponding to another desired receptor segment. The fragments were then combined into a single reaction and PCR amplified a second time with primers corresponding to the 3’ and 5’ ends of the desired chimera and overhangs corresponding to the pCS2+ backbone. The amplified fragments required to generate the chimeric receptor were combined with a linearized pCS2+ backbone and annealed using an In-Fusion kit (Takara Bio, 638947). The resulting plasmids were transformed into Top10 chemically competent cells. Single colonies were submitted for Sanger sequencing to ensure the desired chimera was generated.

### mRNA synthesis

All constructs used in this study were cloned into the pCS2+ expression vector for mRNA synthesis. mRNA was synthesized *in vitro* using the SP6 mMessage mMachine kit (Sigma Aldrich, AM1340) and purified using phenol:chloroform/ethanol extraction. mRNA was stored at −80°C in nuclease free water and aliquoted into 2-4 μl volumes. mRNA concentration was measured using a NanoDrop spectrophotometer. A new aliquot was used for each experiment to prevent mRNA damage from multiple freeze/thaw events.

Due to inconsistent 5’ cap incorporation during mRNA synthesis, mRNA activity could not be projected based on mRNA concentration. Each mRNA injection amount was instead determined individually based on phenotypic evaluation. The following amounts of mRNA were injected into embryos: human intact and GS loop mutant *Acvr1* mRNAs – 100-400 pg, zebrafish *acvr1l* and *acvr1l-R203H* – 100-200 pg, human zebrafish chimeras – 50-200 pg.

### Morpholinos

Morpholinos were synthesized by Gene Tools LLC and stored at −80°C at 25 mg/mL in Danieau solution. Morpholinos were injected at established concentrations for consistent knockdown of their target gene^11^. The following ng injection amounts in 1.3 nl were used for knockdown: endogenous *acvr1l* – mix of 2.3 ng Alk8MO4 and 9.2 ng Alk8MO2; *bmpr1ba* and *bmpr1bb* (*bmpr1b* KD)– mix of 5 ng Alk6aMO1 and 2 ng Alk6bMO1. All morpholinos inhibit translation by binding independent sequences upstream of their respective target gene open reading frame^11,53^.

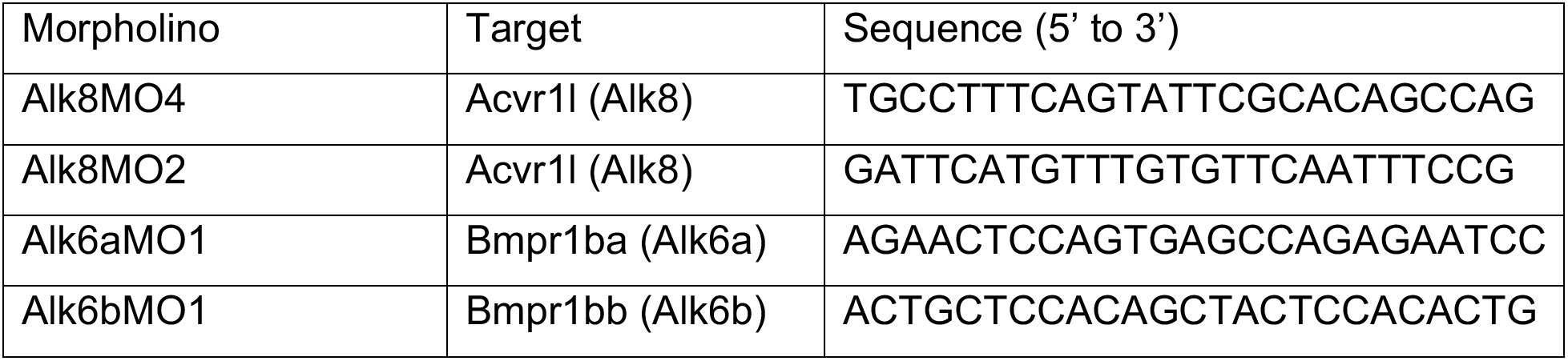

### Injection

mRNA or morpholinos were diluted in nuclease free water or Danieau solution, respectively, with 0.1M KCl and 5-10% phenol red for injection. Eggs at 0-15 min post fertilization were collected and mounted in agarose wells. Embryos were maintained in 22°C E3 media for 15 minutes to slow development during injection and then returned to 28°C. For each experiment an mRNA or morpholino was loaded into a single calibrated needle for injection. Each mRNA or morpholino mix was injected in a volume of 1.3 nl. Needle calibration was checked every 30 min to ensure injection consistency and recalibrated if necessary. For multiple injections into a single embryo, the order in which mRNAs were injected was switched for each plate of embryos. For example, a plate of embryos would be injected with a mRNA, after which a portion of the embryos would be removed as controls, while the remainder of embryos were injected with morpholino. This would then be repeated with the morpholino and mRNA order switched. Embryos from different crosses, injected on different plates on the same day with the same calibrated needles were grouped as a single experiment.

### Phenotyping

Embryos were evaluated for dorsoventral patterning phenotypes at 12-14 hpf (approximately the 8-somite stage) and 28-32 hpf (fully developed body plan and beginning to develop pigment). Embryos were phenotyped based on dorsoventral patterning phenotypes (Fig 1A). Unfertilized embryos were removed at 6-8 hpf. Because C5 embryos lyse by 24 hpf, elongated embryos were separated at 12-14 hpf, and checked for patterning or death at 28 hpf to distinguish between C5 and C4 phenotypes. Bubble plots were generated as previously described by determining the proportion of embryos across multiple experiments that fell into each phenotypic category^25^. Embryo phenotype pictures were obtained using a Leica IC80HD camera and brightfield dissection microscope. Embryos were photographed in E3 media at 28°C at 12-14 hpf and 28-32 hpf. Because mRNA and morpholinos slow the rate of development photographs were taken within the specified time ranges based on embryo stage. Phenotype pictures were white balanced using Adobe photoshop.

### Immunostaining

Embryos were fixed at shield to 60% epiboly stage (approximately 6-7 hpf) in 4% formaldehyde/PBS-Triton. Immunostaining was performed as previously described^25^. Briefly, embryos were blocked in 10% FBS PBS-Triton and probed overnight at 4°C with primary antibody: anti-HA, anti-Flag, or anti-V5, and anti-β-Catenin diluted in blocking solution. Primary antibody was washed, and embryos were probed with secondary antibody for 2 hr at 28°C or overnight at 4°C: anti-Rabbit Alexa 647, anti-Mouse IgG1 Alexa 594, anti-Mouse IgG2a Alexa 594, anti-Mouse Alexa 488, and/or Sytox green diluted in blocking solution. Stained embryos were stored in PBS-Triton at 4°C. No more than one week prior to imaging, embryos were gradually dehydrated in MeOH. For DAPI stained embryos, 1:2000 DAPI was added in a single wash prior to dehydrating embryos in MeOH. Dehydrated embryos were cleared in Benzyl benzoate:Benzyl Alcohol (2:1) no more than 2 hours prior to imaging. Embryos were mounted with the DV axis parallel to the coverslip and imaged using a Zeiss 880 confocal laser microscope. Images were taken at 63x in a 225 x 225 μm frame.

| Antibody | Company/Cat#<br>or RRID | Host<br>species | Application | Dilution | Details |
| --- | --- | --- | --- | --- | --- |
| Anti-HA | RRID:<br>AB_87935 | Rabbit<br>polyclonal | IHC | 1:200 | Diluted in 10%<br>FBS PBS-Triton;<br>Incubated<br>overnight 4°C |
| Anti-Flag | RRID:<br>AB_439687 | Rabbit<br>polyclonal | IHC | 1:500 | Diluted in 10%<br>FBS PBS-Triton;<br>Incubated<br>overnight 4°C |
| Anti-V5 | RRID:<br>AB_2556564 | Mouse<br>IgG2a | IHC | 1:500 | Diluted in 10%<br>FBS PBS-Triton;<br>Incubated<br>overnight 4°C |
| Anti-<br>βCatenin | RRID:<br>AB_476865 | Mouse<br>IgG1 | IHC | 1:1000 | Diluted in 10%<br>FBS PBS-Triton;<br>Incubated<br>overnight 4°C |
| anti-Rabbit<br>Alexa 647 | RRID:<br>AB_2535813 | Goat<br>polyclonal | IHC | 1:500 | Diluted in 10%<br>FBS PBS-Triton;<br>Incubated 2 hr<br>28°C or<br>overnight 4°C |
| anti-Mouse<br>IgG1 Alexa<br>594 | RRID:<br>AB_141592 | Goat<br>polyclonal | IHC | 1:500 | Diluted in 10%<br>FBS PBS-Triton; |
|  |  |  |  |  | Incubated 2 hr<br>28°C or<br>overnight 4°C |
| anti-Mouse<br>IgG2a<br>Alexa 594 | RRID:<br>AB_2535774 |  | IHC | 1:500 | Diluted in 10%<br>FBS PBS-Triton;<br>Incubated 2 hr<br>28°C or<br>overnight 4°C |
| anti-Mouse<br>IgG1 Alexa<br>488 | RRID:<br>AB_2535764 |  | IHC | 1:500 | Diluted in 10%<br>FBS PBS-Triton;<br>Incubated 2 hr<br>28°C or<br>overnight 4°C |
| Sytox<br>green | Fisher/S7020 | NA | IHC | 1:2000 | Diluted in 10%<br>FBS PBS-Triton;<br>Incubated 2 hr<br>28°C or<br>overnight 4°C |
| DAPI | Invitrogen/<br>D3571 | NA | IHC | 1:1000 | Diluted in PBS-Triton 10 min<br>28°C |

