## Supplemental Figures for "Reduced GS domain serine/threonine requirements of Fibrodysplasia Ossificans Progressiva mutant type I BMP receptor ACVR1 in the zebrafish"

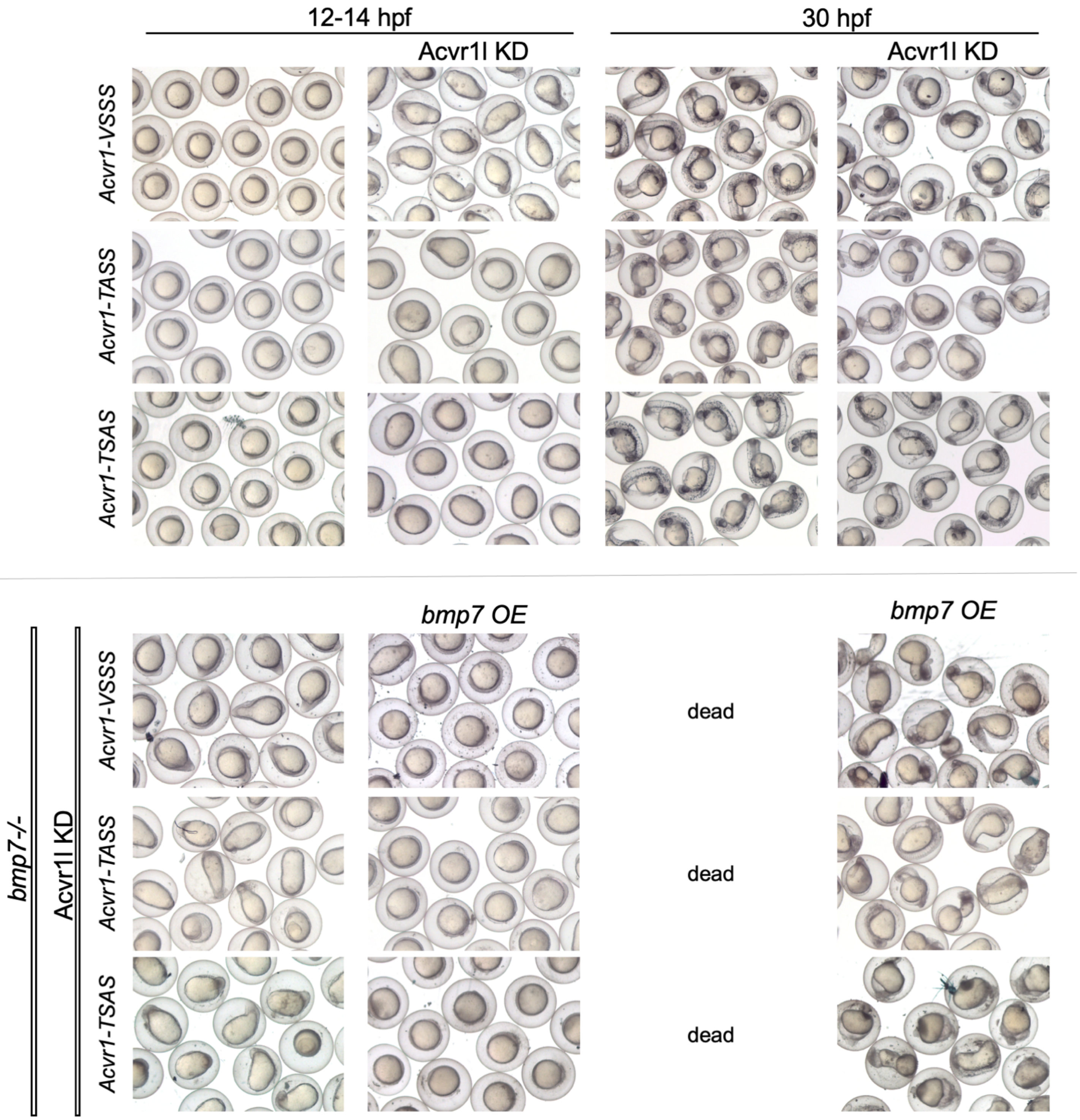


**Supplemental Figure 1:** Representative phenotypes of wild-type and *bmp7-/-* zebrafish embryos with or without Acvr1l KD and/or *bmp7* mRNA OE at 12 and 30 hpf that were injected at the one-cell stage with various *WT-Acvr1* GS loop mutant mRNAs (Fig 2A-A”). Only live embryos are shown at 30 hpf.

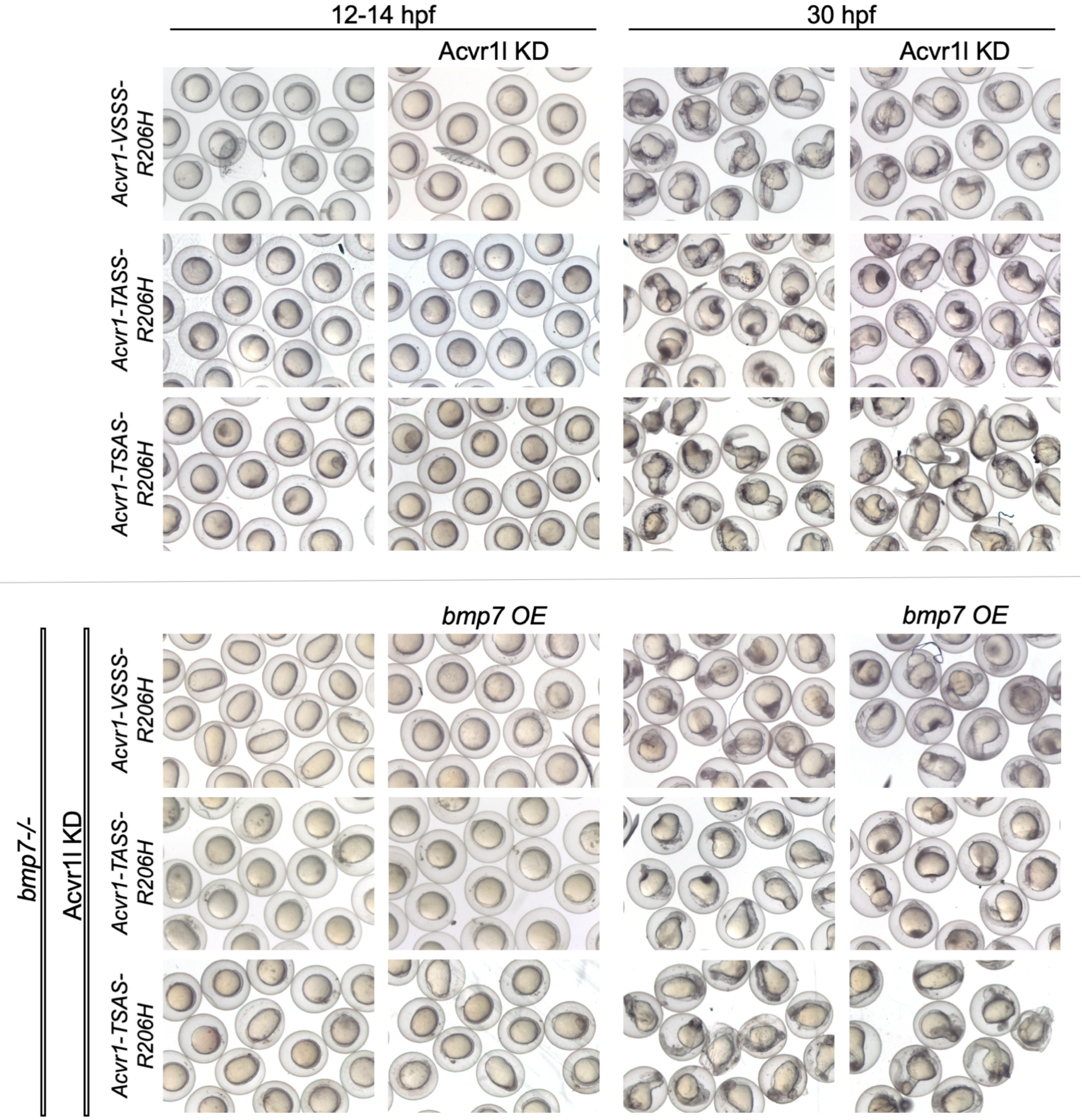


**Supplemental Figure 2** Representative phenotypes of wild-type and *bmp7-/-* zebrafish embryos with or without Acvr1l KD and/or *bmp7* mRNA OE at 12 and 30 hpf that were injected at the one-cell stage with various *Acvr1-R206H* GS loop mutant mRNAs (Fig 2B-B"). Only live embryos are shown at 30 hpf.

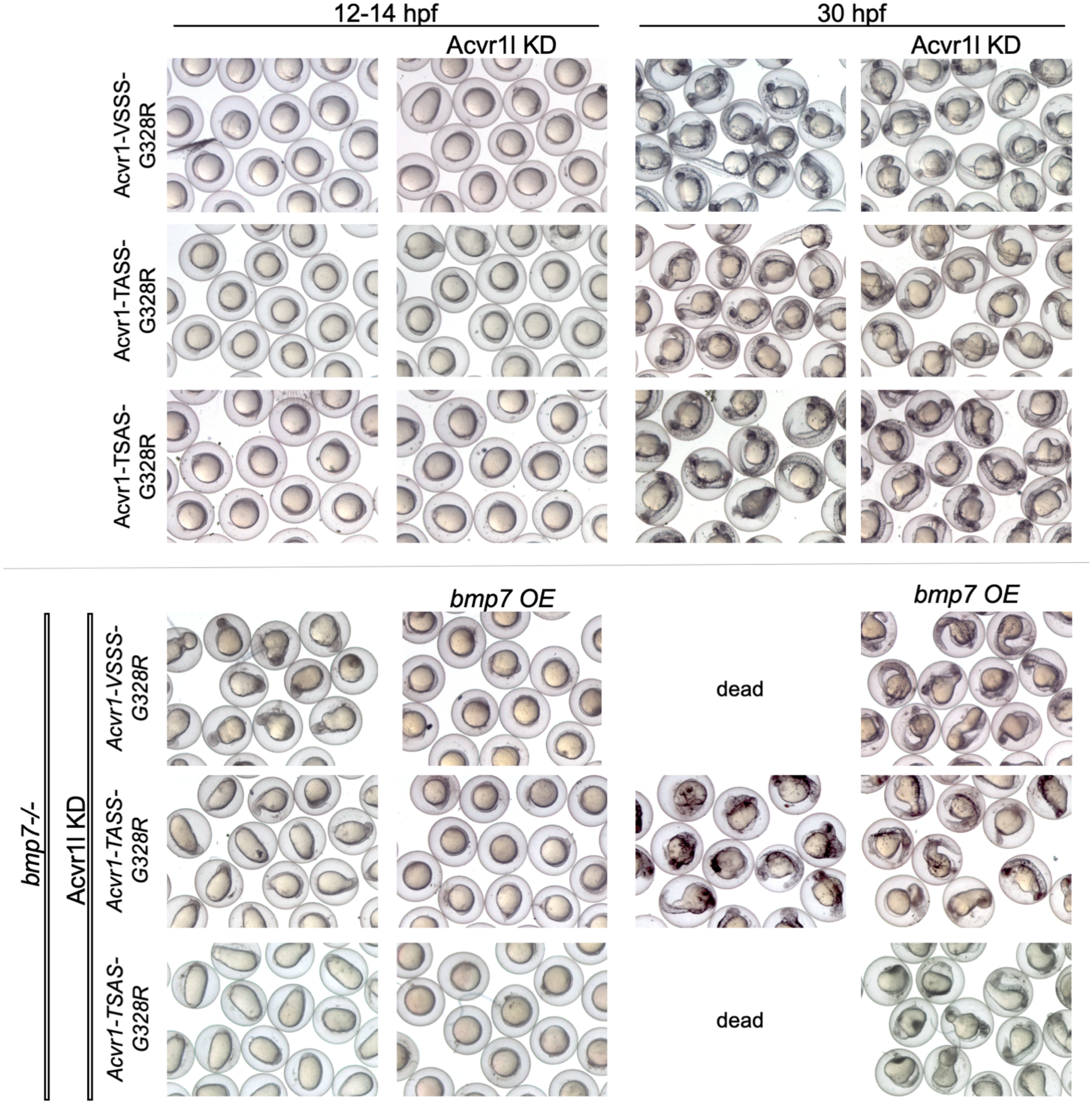


**Supplemental Figure 3:** Representative phenotypes of wild-type and *bmp7-/-* zebrafish embryos with or without Acvr1l KD and/or *bmp7* mRNA OE at 12 and 30 hpf that were injected at the one-cell stage with various *Acvr1-G328R* GS loop mutant mRNAs (Fig 2C-C"). Only live embryos are shown at 30 hpf.

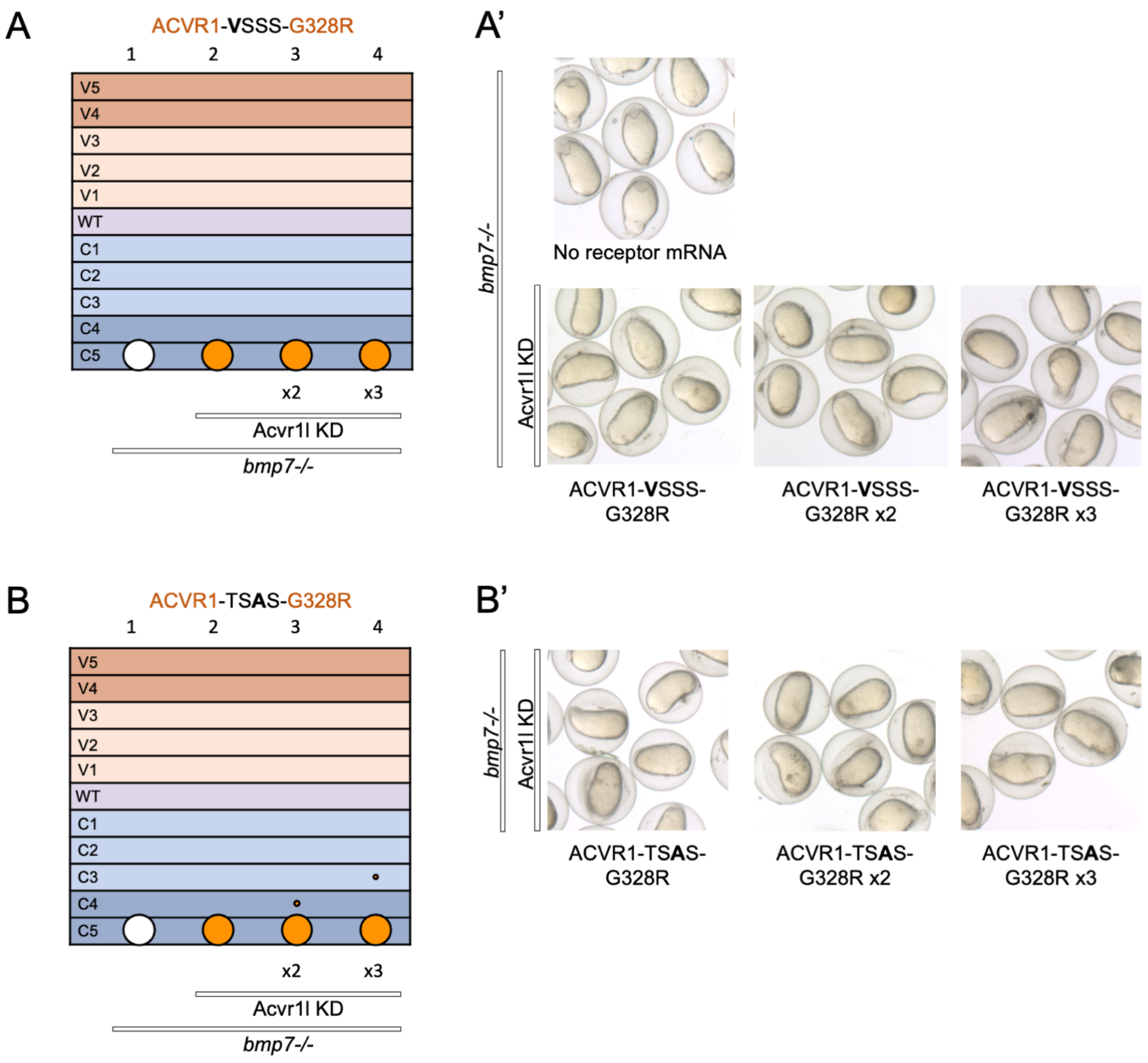


**Supplemental Figure 4** *Acvr1-G328R* GS loop mutant mRNA titration in *bmp7-/-* embryos with Acvr1l KD. Receptor mRNA was injected at normal, double (x2) or triple (x3) amounts. **(A)** Acvr1-**V**SSS-G328R, column: 1, N=118; 2, N=42; 3, N=38; 4, N=31. **(A')** Representative 12 hpf phenotypes. **(B)** Acvr1-TS**A**S-G328R, column: 1, N=118; 2, N=70; 3, N=55; 4 N=43. **(B')** Representative 12 hpf phenotypes. 28-30 hpf phenotypes not shown because 95% of embryos were dead by this time. Injections were performed over two experiments. Supplemental Fig 4A and 4B share control uninjected embryos.

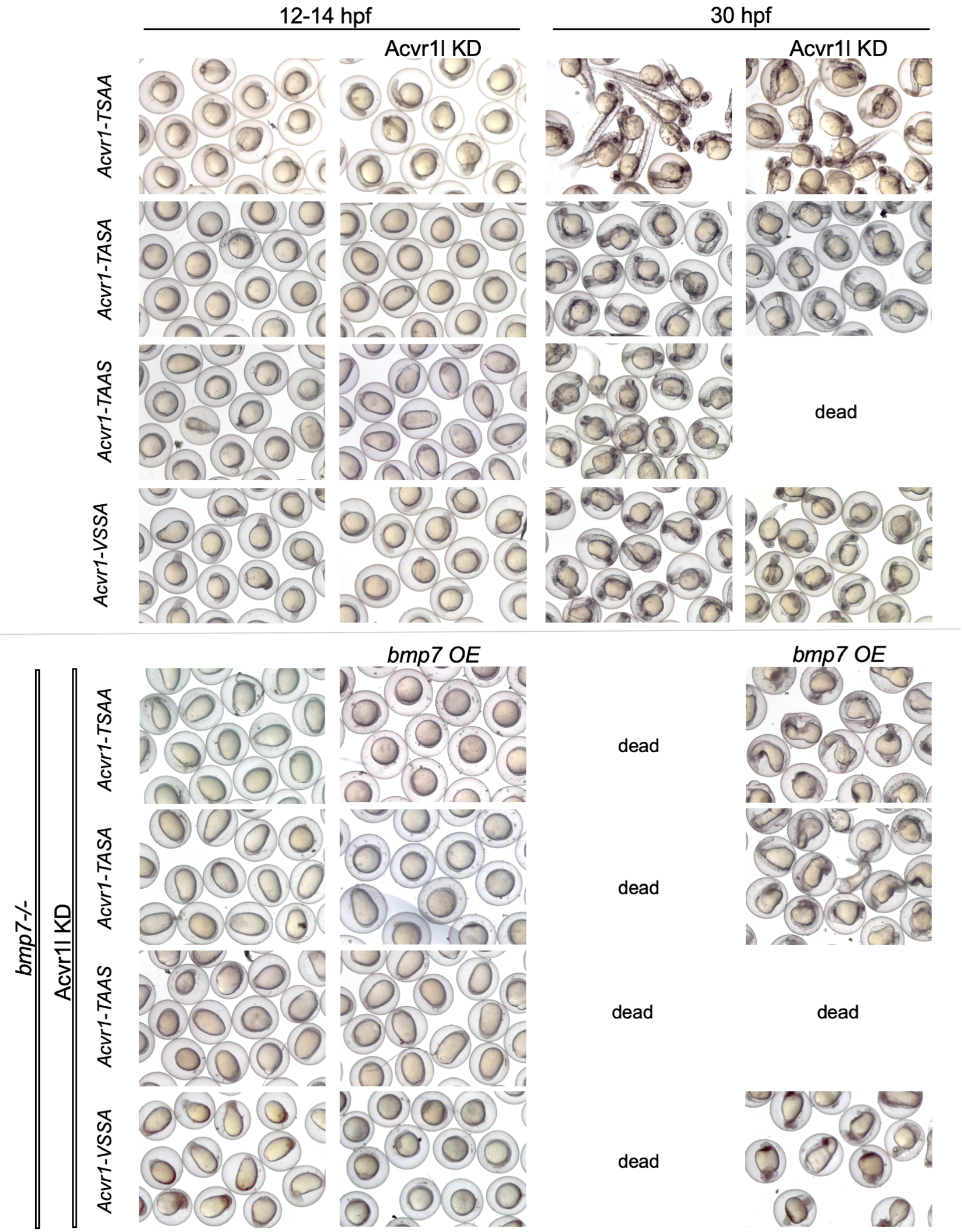

12-14 hpf
30 hpf
Acvr1l KD
Acvr1l KD
*Acvr1-TSAA*
*Acvr1-TASA*
*Acvr1-TAAS*
*Acvr1-VSSA*
dead
*bmp7 OE*
*bmp7 OE*
*bmp7-/-*
Acvr1l KD
*Acvr1-TSAA*
*Acvr1-TASA*
*Acvr1-TAAS*
*Acvr1-VSSA*
dead
dead
dead
dead
dead

**Supplemental Figure 5:** Representative phenotypes of wild-type and *bmp7-/-* zebrafish embryos with or without Acvr1l KD and/or *bmp7* mRNA OE at 12 and 30 hpf that were injected at the one-cell stage with various *WT-Acvr1* GS loop mutant mRNAs (Fig 3A-A’’’). Only live embryos are shown at 30 hpf.

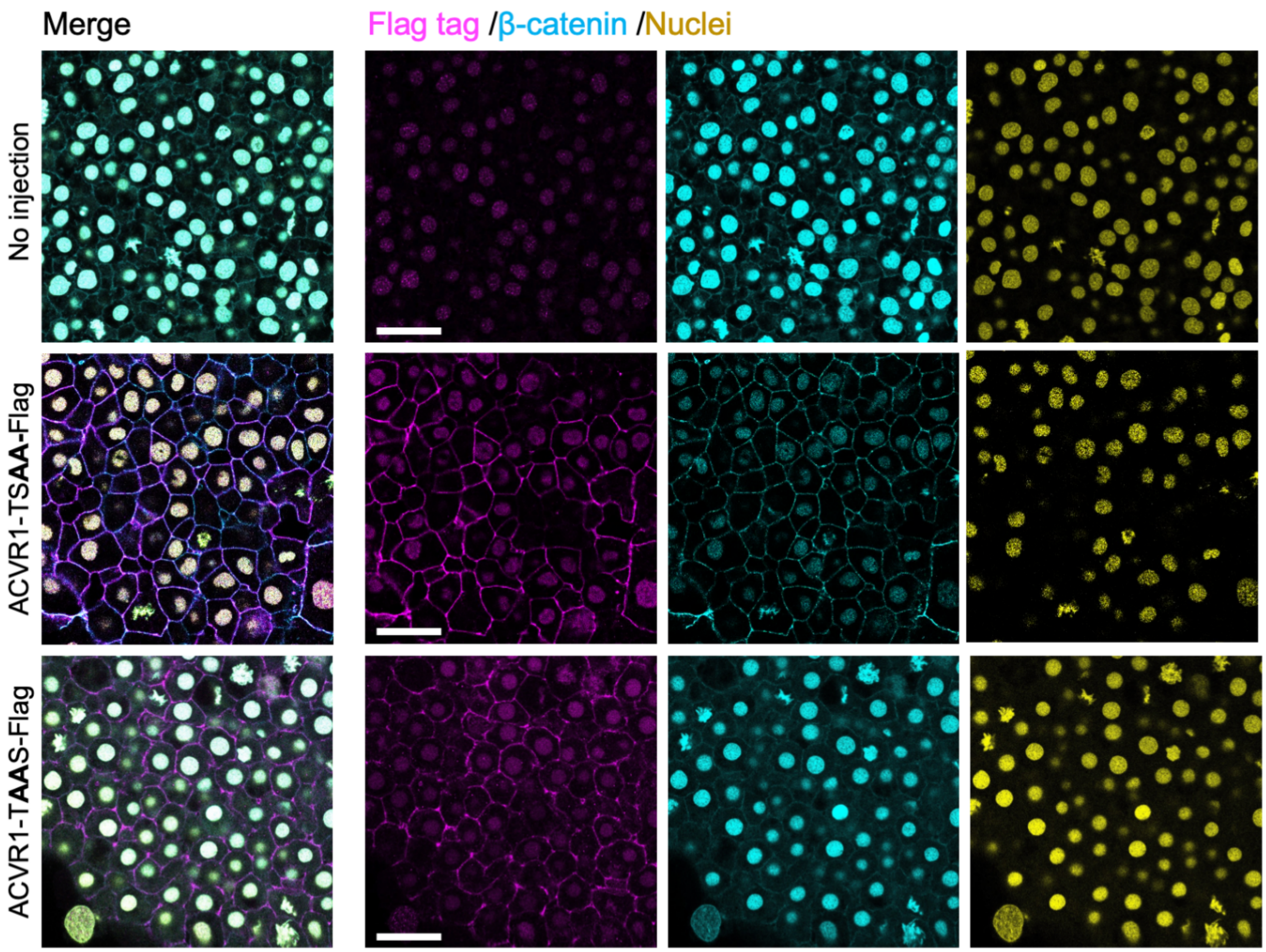


**Supplemental Figure 6:** Nuclei, β-Catenin and Flag immunostaining in 12 hpf zebrafish embryos injected with mRNAs for Flag-tagged GS loop mutant WT- or FOP-ACVR1. Scale bars = 25 μm. No injection (N=6), ACVR1-TSAA-Flag (N=6), ACVR1-TAAS-Flag (N=6).

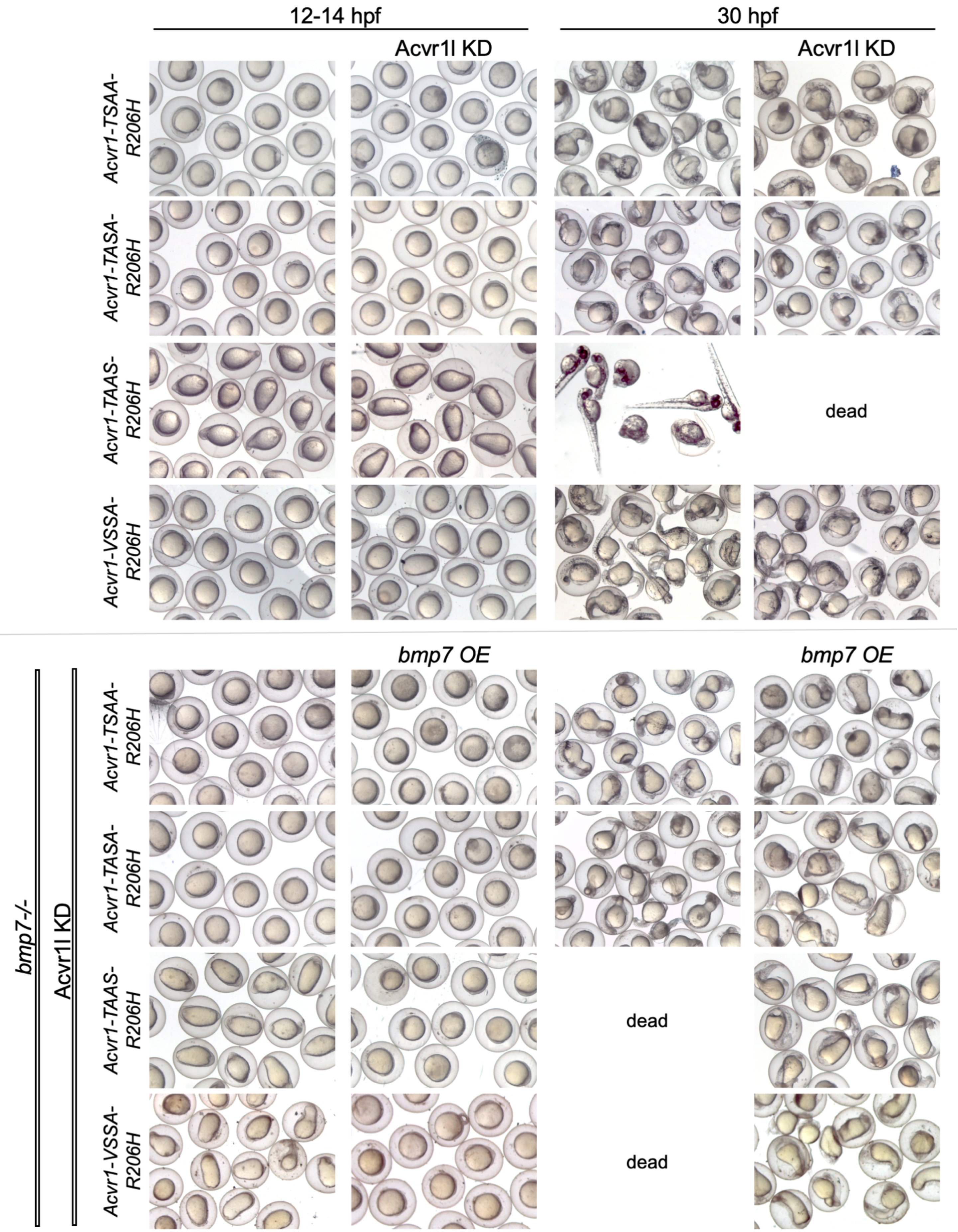
12-14 hpf
30 hpf
Acvr1l KD
Acvr1l KD
Acvr1-TSAA-R206H
Acvr1-TASA-R206H
Acvr1-TAAS-R206H
Acvr1-VSSA-R206H
dead
bmp7 OE
bmp7 OE
bmp7-/-
Acvr1l KD
Acvr1-TSAA-R206H
Acvr1-TASA-R206H
Acvr1-TAAS-R206H
Acvr1-VSSA-R206H
dead
dead

**Supplemental Figure 7:** Representative phenotypes of wild-type and *bmp7-/-* zebrafish embryos with or without Acvr1l KD and/or *bmp7* mRNA OE at 12 and 30 hpf that were injected at the one-cell stage with various *Acvr1-R206H* GS loop mutant mRNAs (Fig 3B-B’’’). Only live embryos are shown at 30 hpf.

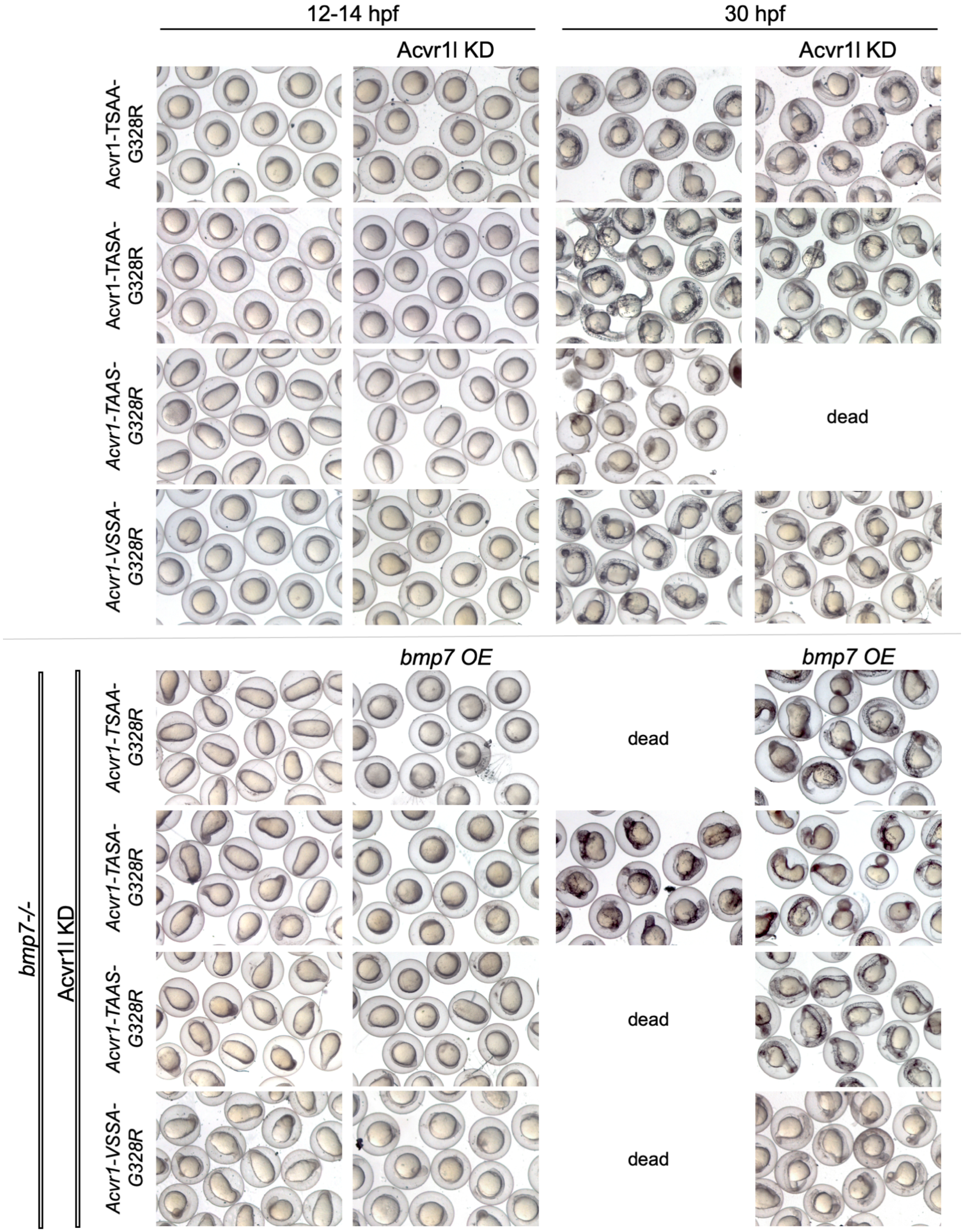

12-14 hpf
30 hpf
Acvr1l KD
Acvr1l KD
Acvr1-TSAA-G328R
Acvr1-TASA-G328R
Acvr1-TAAS-G328R
Acvr1-VSSA-G328R
dead
bmp7 OE
bmp7 OE
bmp7-/-
Acvr1l KD
Acvr1-TSAA-G328R
Acvr1-TASA-G328R
Acvr1-TAAS-G328R
Acvr1-VSSA-G328R
dead
dead
dead

**Supplemental Figure 8:** Representative phenotypes of wild-type and *bmp7-/-* zebrafish embryos with or without Acvr1l KD and/or *bmp7* mRNA OE at 12 and 30 hpf that were injected at the one-cell stage with various *Acvr1-G328R* GS loop mutant mRNAs (Fig 3C-C'''). Only live embryos are shown at 30 hpf.

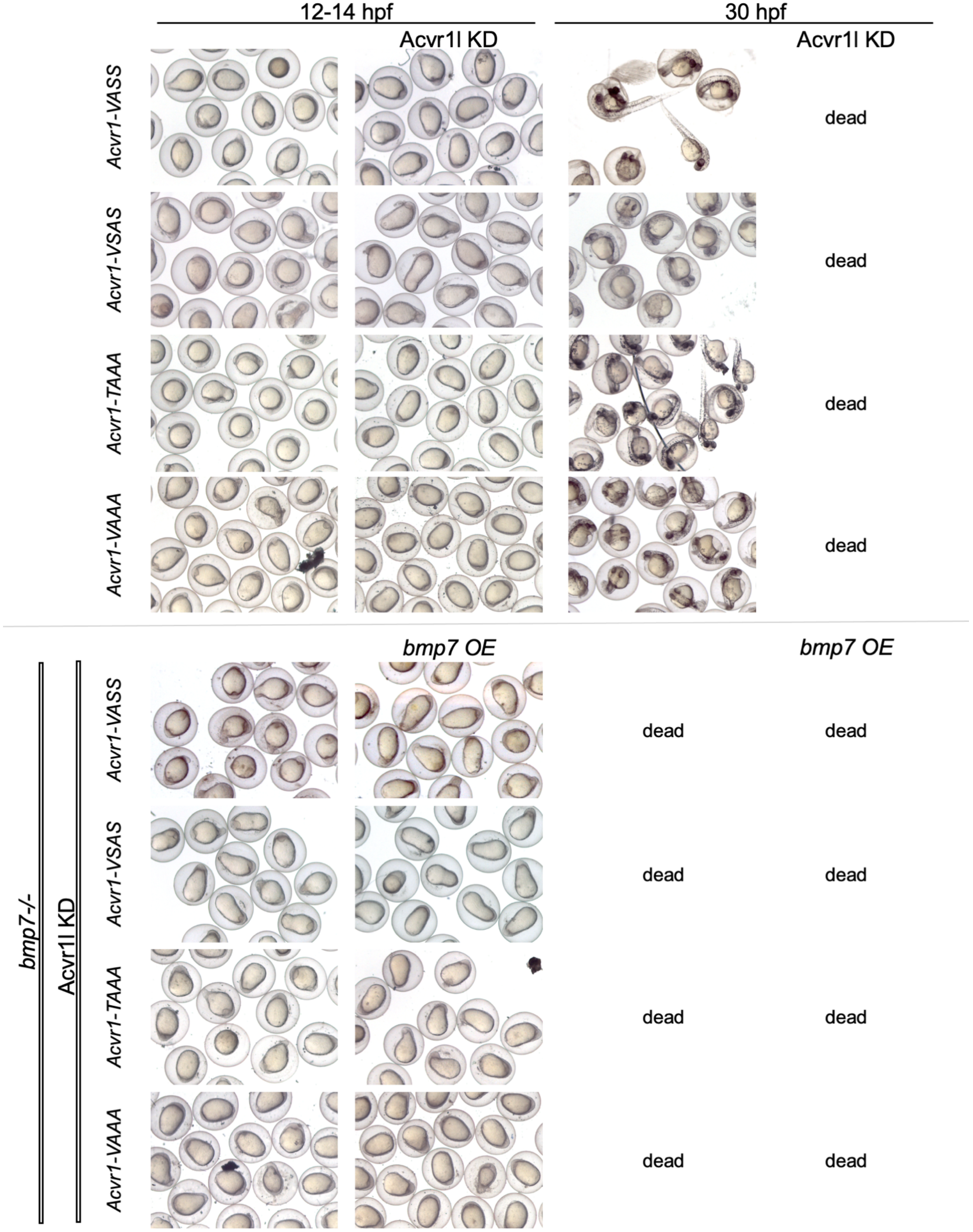

12-14 hpf
30 hpf
Acvr1l KD
Acvr1l KD
Acvr1-VASS
dead
Acvr1-VSAS
dead
Acvr1-TAAA
dead
Acvr1-VAAA
dead
bmp7 OE
bmp7 OE
bmp7-/-
Acvr1l KD
Acvr1-VASS
dead
dead
Acvr1-VSAS
dead
dead
Acvr1-TAAA
dead
dead
Acvr1-VAAA
dead
dead

**Supplemental Figure 9:** Representative phenotypes of wild-type and *bmp7-/-* zebrafish embryos with or without Acvr1l KD and/or *bmp7* mRNA OE at 12 and 30 hpf that were injected at the one-cell stage with various *WT-Acvr1* GS loop mutant mRNAs (Fig 4A-A’’’). Only live embryos are shown at 30 hpf.

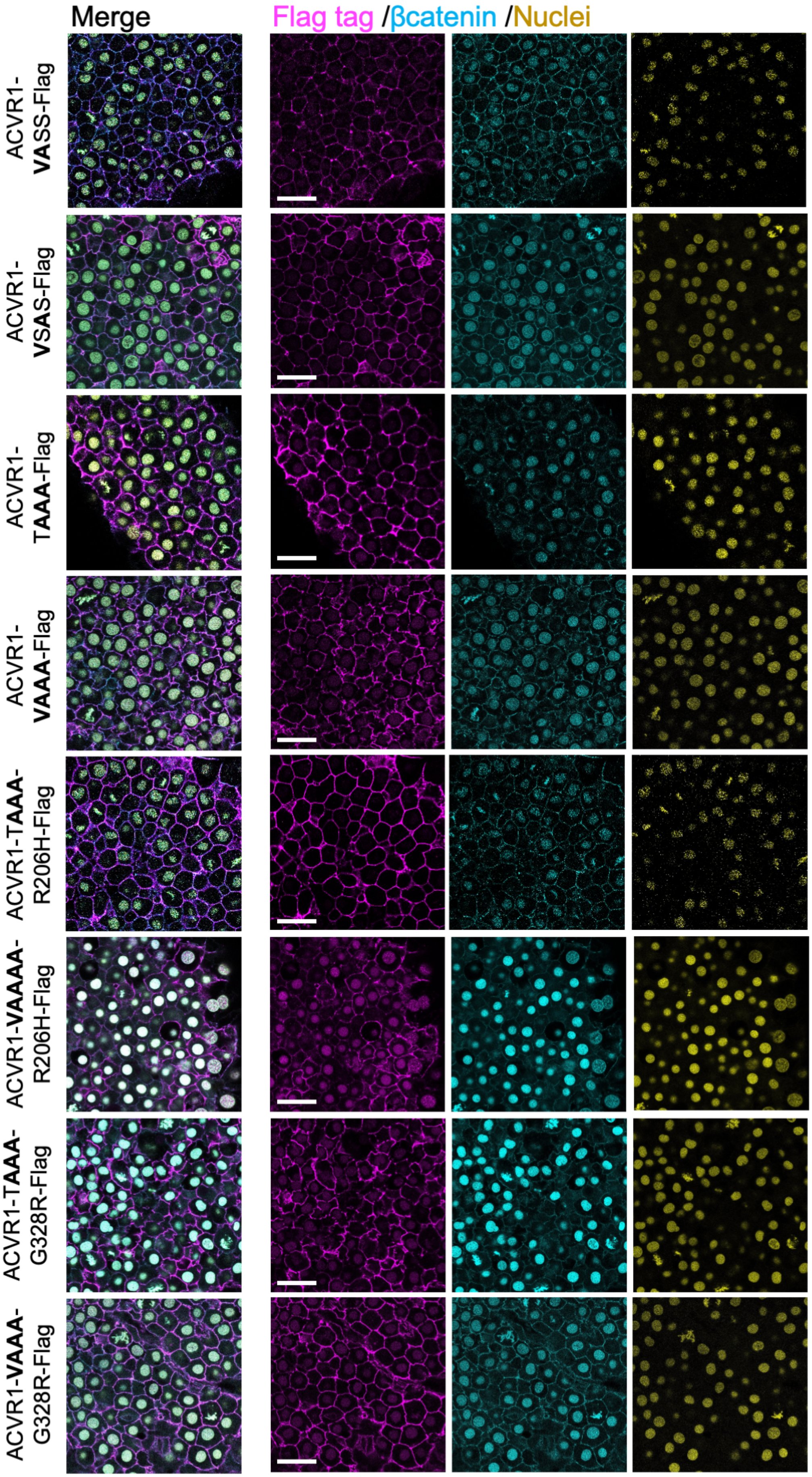

Merge
Flag tag /βcatenin /Nuclei
ACVR1-VASS-Flag
ACVR1-VSAS-Flag
ACVR1-TAAA-Flag
ACVR1-VAAA-Flag
ACVR1-TAAA-R206H-Flag
ACVR1-VAAAA-R206H-Flag
ACVR1-TAAA-G328R-Flag
ACVR1-VAAA-G328R-Flag

**Supplemental Figure 10:** Nuclei, Beta-catenin and Flag immunostaining in 12 hpf zebrafish embryos injected with mRNAs of Flag-tagged GS loop mutant WT- or FOP-ACVR1. Scale bars = 25µm. ACVR1-VASS-Flag (N=5), ACVR1-VSAS-Flag (N=3), ACVR1-TAAA-Flag (N=4), ACVR1-VAAA-Flag (N=5), ACVR1-TAAA-R206H-Flag (N=3), ACVR1-VAAA-R206H-Flag (N=5), ACVR1-TAAA-G328R-Flag (N=3), ACVR1-VAAA-G328R-Flag (N=2).

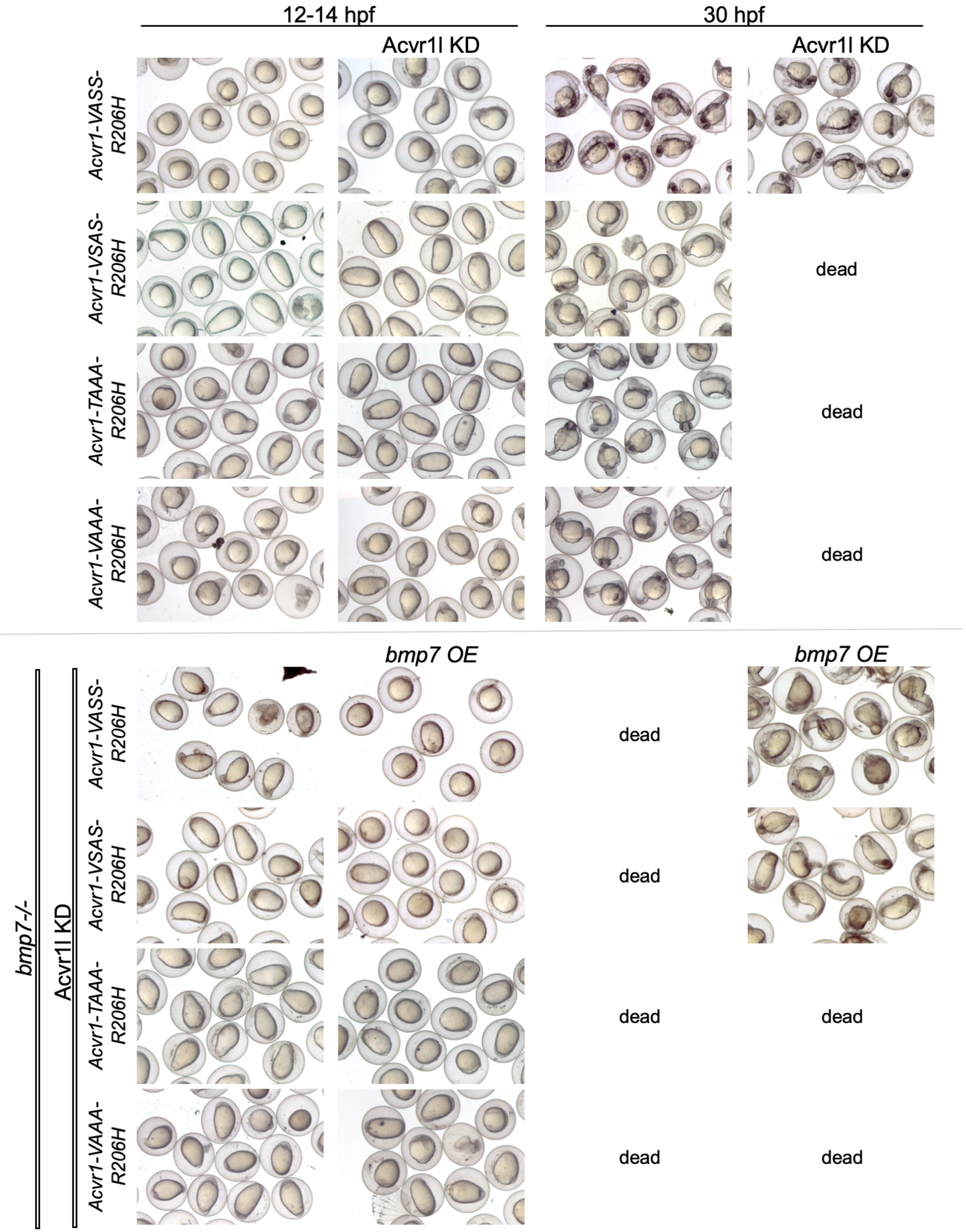
12-14 hpf
30 hpf
Acvr1l KD
Acvr1l KD
Acvr1-VASS-R206H
Acvr1-VSAS-R206H
Acvr1-TAAA-R206H
Acvr1-VAAA-R206H
dead
dead
dead
bmp7 OE
bmp7 OE
bmp7-/-
Acvr1l KD
Acvr1-VASS-R206H
Acvr1-VSAS-R206H
Acvr1-TAAA-R206H
Acvr1-VAAA-R206H
dead
dead
dead
dead
dead
dead

**Supplemental Figure 11:** Representative phenotypes of wild-type and *bmp7-/-* zebrafish embryos with or without Acvr1l KD and/or *bmp7* mRNA OE at 12 and 30 hpf that were injected at the one-cell stage with various *Acvr1-R206H* GS loop mutant mRNAs (Fig 4B-B’’’). Only live embryos are shown at 30 hpf.

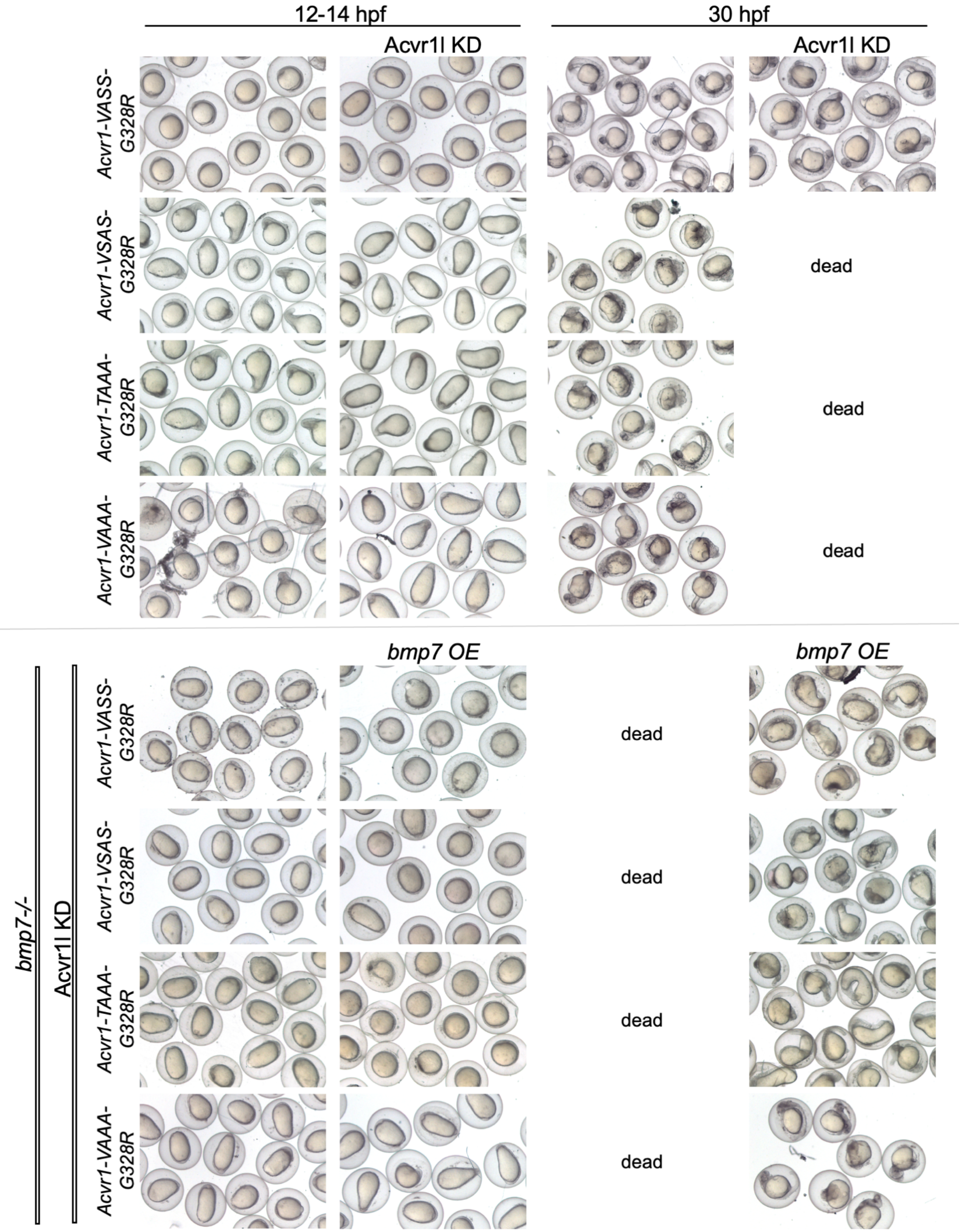
12-14 hpf
30 hpf
Acvr1l KD
Acvr1l KD
Acvr1-VASS-G328R
Acvr1-VSAS-G328R
dead
Acvr1-TAAA-G328R
dead
Acvr1-VAAA-G328R
dead
bmp7 OE
bmp7 OE
bmp7-/-
Acvr1l KD
Acvr1-VASS-G328R
dead
Acvr1-VSAS-G328R
dead
Acvr1-TAAA-G328R
dead
Acvr1-VAAA-G328R
dead

**Supplemental Figure 12:** Representative phenotypes of wild-type and *bmp7-/-* zebrafish embryos with or without Acvr1l KD and/or *bmp7* mRNA OE at 12 and 30 hpf that were injected at the one-cell stage with various *Acvr1-G328R* GS loop mutant mRNAs (Fig 4C-C’’’). Only live embryos are shown at 30 hpf.

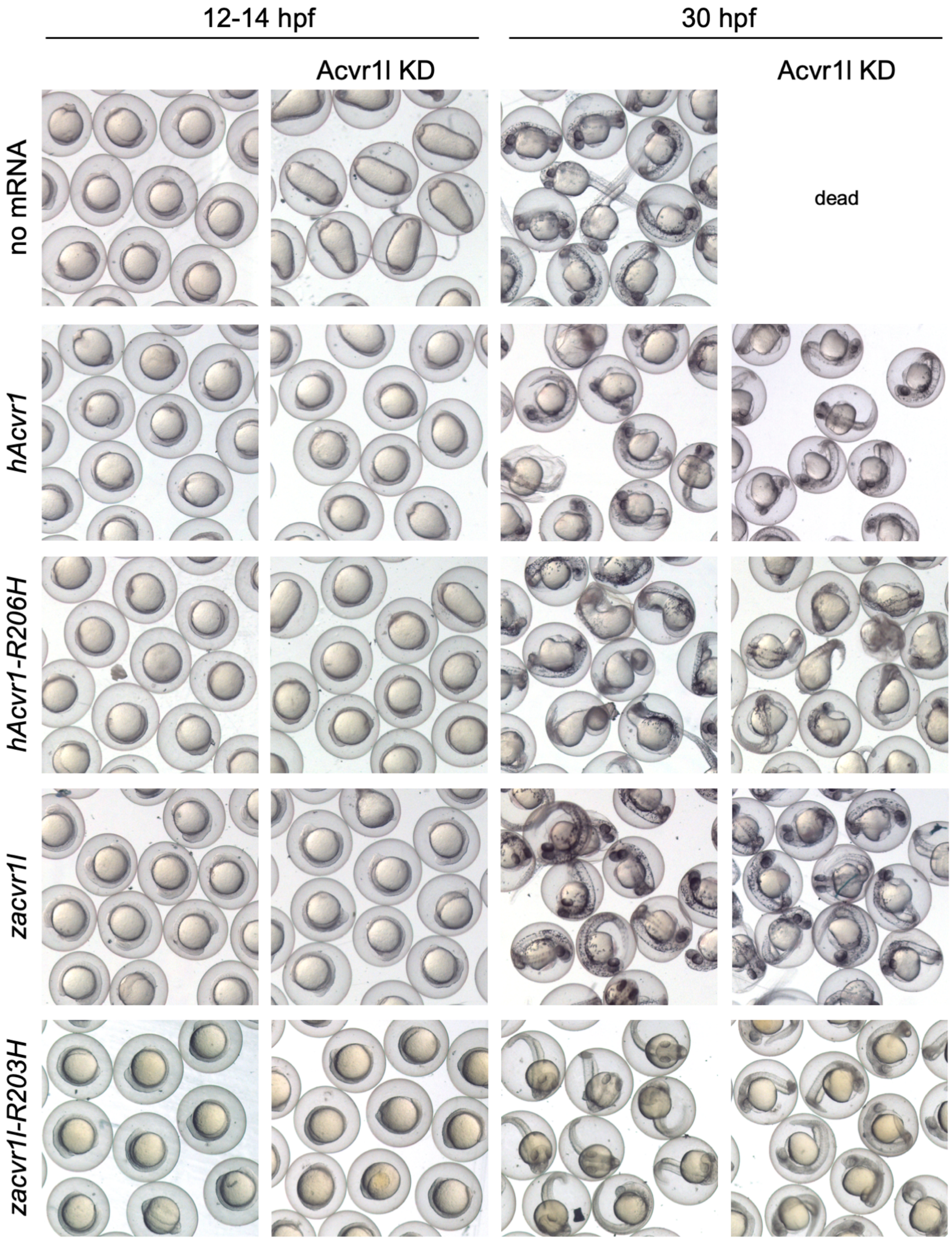
12-14 hpf
30 hpf
Acvr1l KD
Acvr1l KD
no mRNA
dead
*hAcvr1*
*hAcvr1-R206H*
*zacvr1l*
*zacvr1l-R203H*

**Supplemental Figure 13:** Representative 12 and 24 hpf phenotypes of wild-type embryos with or without Acvr1l KD injected with human (h)*Acvr1*, *hAcvr1-R206H*, zebrafish (z)*acvrl1* or *zacvr1l-R203H* mRNA (Fig 6C). Only live embryos are shown at 30 hpf.

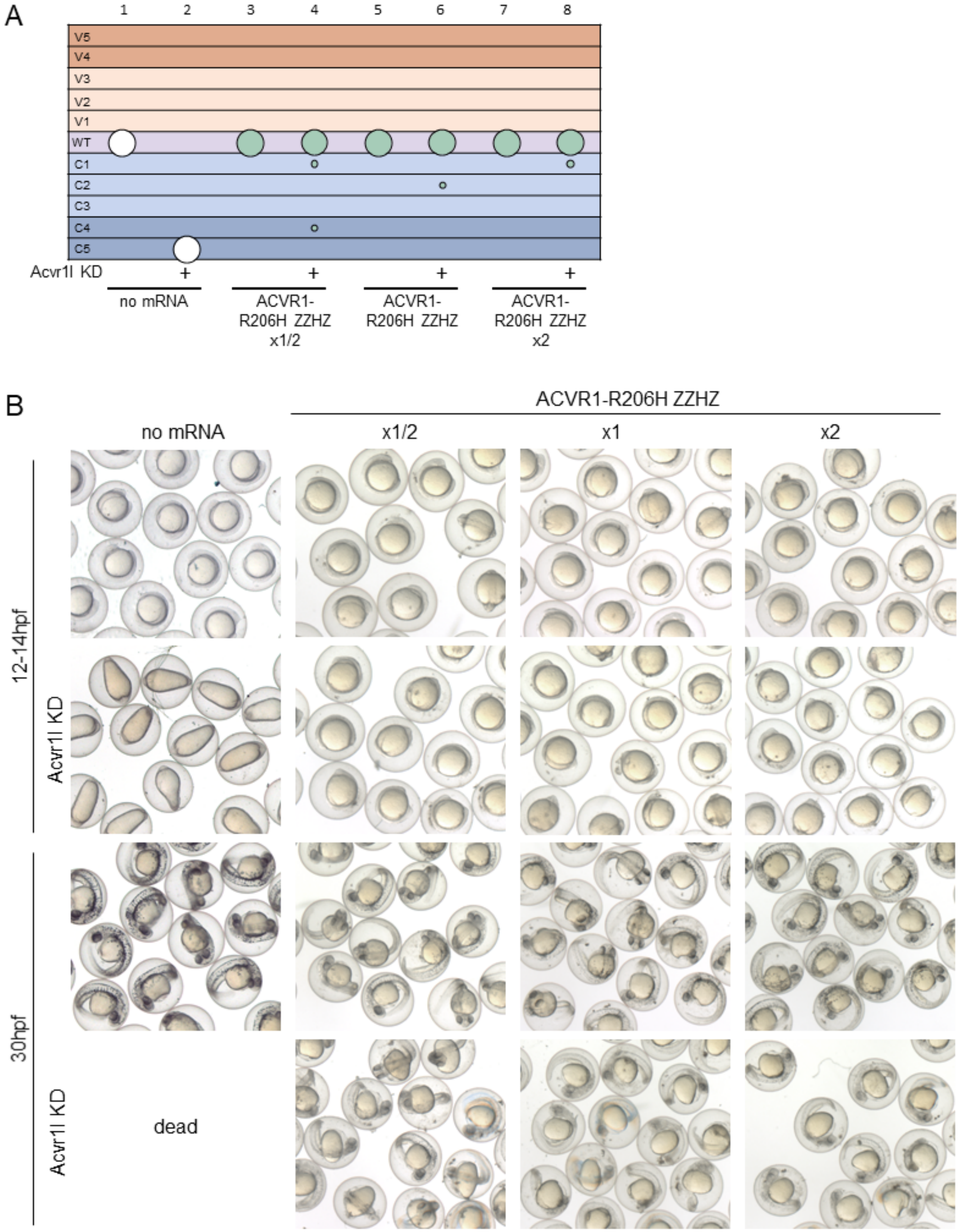


**Supplemental Figure 14:** *Acvr1-R206H-ZZHZ* titration **(A)** *Acvr1-R206H-ZZHZ* mRNA at half (x1/2), single (x1) or double (x2) amounts injected into wild-type zebrafish embryos with or without *acvr1l* KD. Columns: 1, N=35; 2, N=99; 3, N=43; 4, N=38; 5, N=55; 6, N=60; 7, N=38; 8, N=47. **(B)** Representative

12 and 30 hpf phenotypes. Injection was performed over two experiments. Only live embryos are shown at 30 hpf (B).

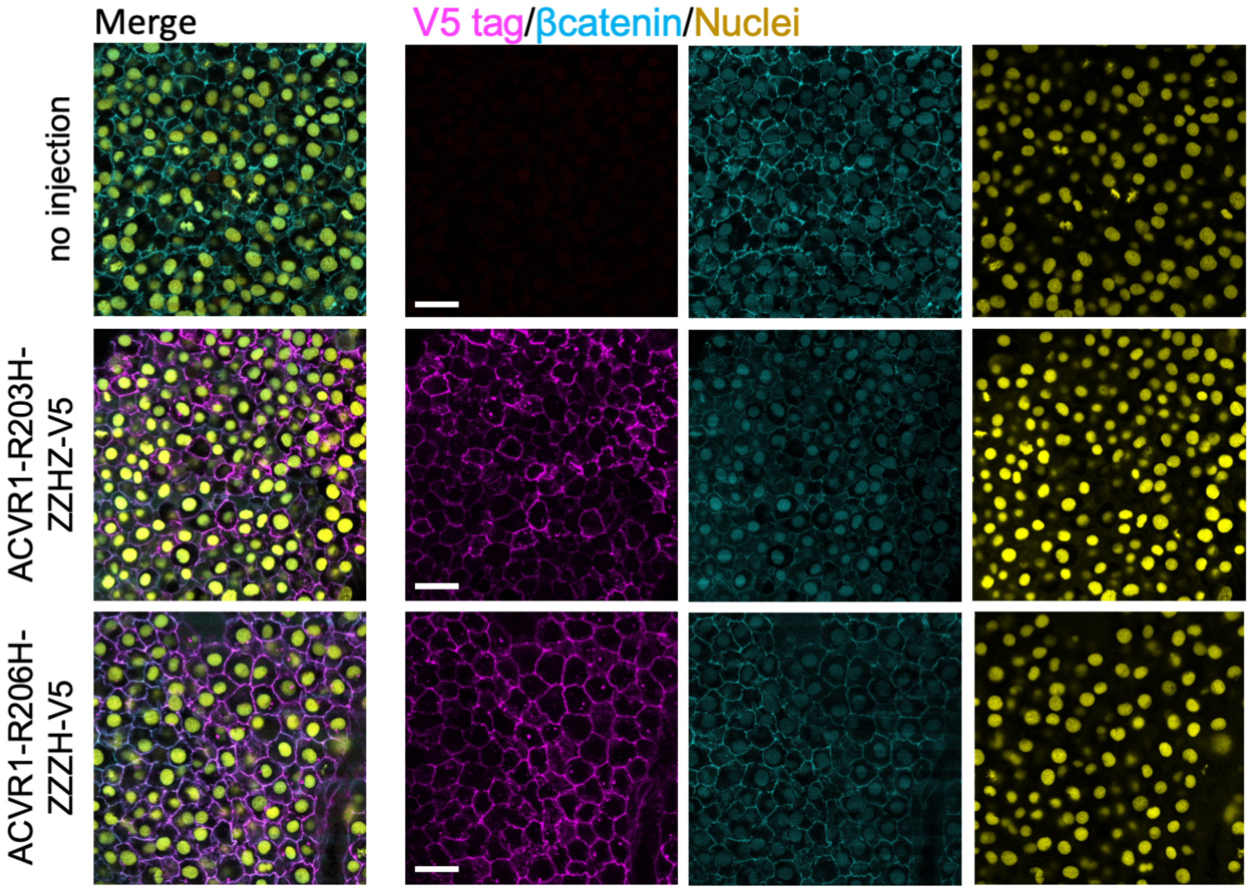


**Supplemental Figure 15:** Nuclei, β-catenin and V5 immunostains in 12 hpf zebrafish embryos injected with mRNA for V5-tagged chimeric zebrafish-human ACVR1. No injection (N=3), ACVR1-R203H-ZZHZ (N=5), ACVR1-R206H-ZZZH (N=4).

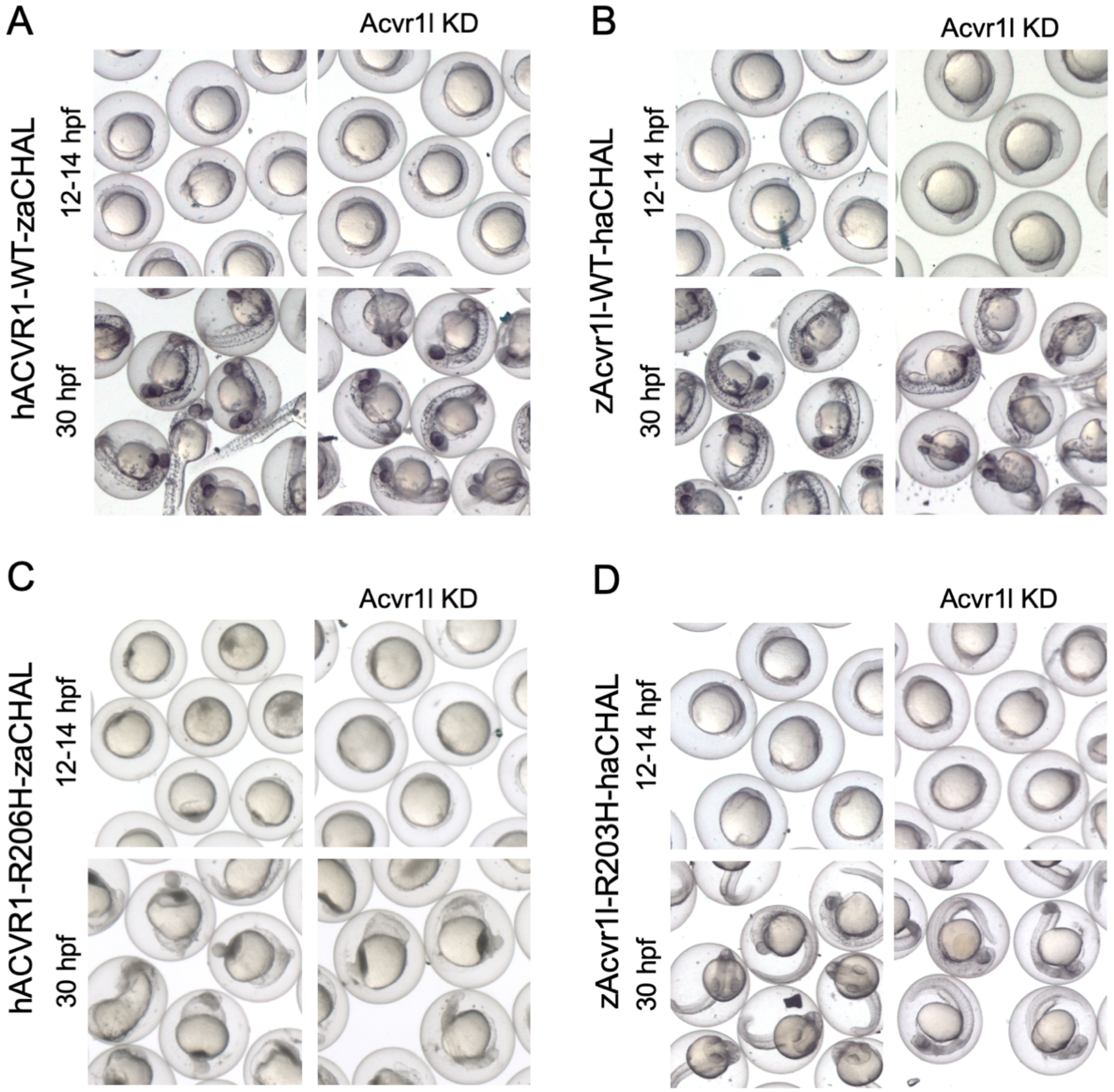


**Supplemental Figure 16:** Representative 12 and 24 hpf phenotypes of wild-type embryos with or without Acvr1l KD injected with **(A)** human (h)ACVR1-WT-zaCHAL, **(B)** zebrafish (z)Acvr1l-WT-haCHAL, **(C)** hACVR1-R206H-zaCHAL, or **(D)** zAcvr1l-R203H-haCHAL (Fig. 6G-J). Only live embryos are shown at 30 hpf.
